# Dynamics of peripheral blood flow across sleep stages

**DOI:** 10.1101/2021.11.04.467081

**Authors:** Zhiwei Fan, Yoko Suzuki, Like Jiang, Satomi Okabe, Shintaro Honda, Junki Endo, Takahiro Watanabe, Takashi Abe

## Abstract

**Study Objectives:** Pulse rate variability (PRV) derived from peripheral blood flow has been reported as a surrogate parameter for heart rate variability (HRV). However, there are currently no studies reporting systematic comparisons of PRV with HRV in a normal sleep state. Whether PRV can provide similar information regarding sleep stages remains unclear. Peripheral blood flow may also be modulated differently across sleep stages. Thus, we aimed to investigate blood flow dynamics and compare PRV with HRV across sleep stages to see if blood flow can provide further information about sleep stages.

**Methods:** We performed electrocardiography and simultaneously measured blood flow from the right index finger and ear concha of 45 healthy participants (13 women; mean age, 22.5 ± 3.4 years) during one night of sleep. Time-domain, frequency-domain, and non-linear indices of PRV/HRV, and time- and frequency-domain blood flow parameters were calculated.

**Results:** Finger-PRV results showed similar patterns to HRV results for most parameters. Finger-blood flow parameters in the time and frequency domains also showed information about the different sleep stages. Further, both finger- and ear-blood flow results showed 0.2–0.3 Hz oscillations that varied with sleep stages, with a significant increase in N3, suggesting a modulation (of respiration) within this frequency band.

**Conclusions:** These results suggest that PRV can provide as much information as HRV for different sleep stages. Furthermore, the results show that blood flow + PRV could be more advantageous than HRV alone in the assessment of the sleep state and related autonomic nervous activity.

**Statement of Significance:** The research provides comprehensive information on peripheral blood flow (BF) activity across sleep stages for the first time, as the major novelty of our work. The second contribution is a systematic study comparing BF-derived pulse rate variability (PRV) with heart rate variability across sleep stages in a normal sleep state. We believe that our work makes a significant contribution to the literature because it provides comprehensive information on the potential of BF+PRV as a new biomarker for assessing the sleep state. Further, this study contributes to developing a more convenient method of assessing the sleep state in the clinical and home/work setting.

## 1 Introduction

In sleep medicine, human sleep is often analyzed in sleep-wake stages according to the American Academy of Sleep Medicine (AASM) criteria: wakefulness (Wk), the three stages of non-rapid-eye-movement (NREM) sleep (N1, N2, and N3), and rapid-eye-movement (REM) sleep.[1] Different sleep stages, such as NREM and REM, have very different roles in daily functioning, such as learning and memory.[2] In addition, different sleep stages correspond to different central nervous system (CNS) activity patterns, measured using an electroencephalogram (EEG). In contrast, autonomic nervous system (ANS) activity varies across EEG-defined sleep stages because of CNS-ANS coupling, reflected in heart rate activity and heart rate variability (HRV).[3]

ANS activity can be measured using HRV derived from an electrocardiogram (ECG).[3,4] Pulse rate variability (PRV) is reported to be a surrogate parameter for HRV, which can be measured more conveniently. It is derived from the signals recorded from peripheral sites, such as fingers and ears, using techniques such as photoplethysmography and laser Doppler flowmetry (LDF).[5–7] The validity of PRV as a substitute for HRV depends on different conditions, such as physiological states,[8] posture, environment, and technical factors. Studies have shown that PRV is consistent with HRV during resting states,[9,10] such as sleep. Attempts have been made in previous studies to determine the level of agreement between PRV and HRV during each sleep stage.[11,12] However, these studies focused only on patients with sleep apnea. In addition, they did not involve the comparison of sleep stages with respect to HRV and PRV parameters. For example, although HRV power is found to be modulated by sleep stages[13,14] previous studies did not include quantitative investigations of the differences in HRV power between specific sleep stages. At present, no study has been conducted to systematically compare PRV with HRV with respect to information provided for differentiating stages of normal sleep. Thus, one of the aims of this study was to confirm whether PRV can provide as much information as HRV for the differentiation of sleep stages. Considering the relationship between PRV and HRV revealed in previous studies,[3,8] it is expected that PRV parameters may differ across sleep stages as HRV parameters do. What matters is the information PRV can provide in the sleep state; for example, whether PRV can be favorably compared with HRV in differentiating sleep stages. Recent studies have shown PRV as a new biomarker rather than a surrogate for HRV.15] Thus, there must be similarities and differences between PRV and HRV metrics.

Blood flow (BF) signal may provide further information about sleep stages. Thus, the other purpose of this study was to investigate peripheral BF signals in the time and frequency domains during different sleep stages. To the best of our knowledge, this is the first study to conduct such an investigation. Most previous studies only focused on PRV. ANS activity and related processes during sleep may affect BF and ECG signals differently. The two components of the ANS, the parasympathetic and the sympathetic, have different roles in the cardiovascular system. While the parasympathetic nervous system majorly contributes to heartbeat activity,[16] the sympathetic nervous system plays a role in heartbeat activity and a dominant role in regulating vascular activity.[17,18] Therefore, BF may be affected by factors that also affect an ECG through the blood vessel network from the heart, and by factors that affect vascular activity alone, such as local (metabolic, myogenic, and paracrine) controls.[19] Thus, some of the variabilities in BF signals may come from sources that do not affect ECG signals in the same way. In addition, BF signals may vary even more than ECG signals because of the modulation of respiration.[11,12,20] For example, obstructive respiratory events in sleep apnea lead to more changes in the pulse wave amplitude of BF than in the heart rate derived from an ECG. This shows the effect of respiration on peripheral vascular activity. It also suggests that parameters of BF signals other than PRV indices may matter. Furthermore, respiration-modulated BF oscillations in the peripheral vascular bed are also affected by the sympathetic nervous system through vasomotion.[7] Thus, BF may have distinct characteristics during sleep and is worth investigating. Investigating the dynamics of BF may reveal its potential for determining and predicting neural pattern-defined sleep stages. Therefore, in addition to comparing PRV metrics with those of HRV, we also aimed to investigate the dynamics of BF across different sleep stages to clarify the roles of PRV and BF in monitoring the sleep state.

## 2 Methods

### 2.1 Participants

Forty-five healthy adults (13 women; mean age, 22.5 ± 3.4 years) were recruited and screened for inclusion in this study. All participants were physically and psychologically healthy and were not being treated for any sleep disorder. The included participants satisfied the following criteria: 1) older than 20 years old and younger than 60 years old; 2) able to fill out the Japanese instruction documents, consent forms, and survey forms; 3) able to sleep in the examination rooms in the sleep lab; and 4) not currently being treated for a sleep disorder. Participants were excluded if they were claustrophobic; had uncontrolled diabetes; had a history of myocardial infarction; had unstable angina, serious liver disease, or serious renal disease; were pregnant or may become pregnant; were lactating; or were judged by the investigator to be inappropriate as participants. The Ethics Committees of the University of Tsukuba (ID: R01-101) approved the research protocol. All participants provided written consent and received payment for participation.

### 2.2 Apparatus and procedure

All participants underwent an 8-hour whole-night sleep in a sound-proof chamber and their physiological activities were recorded while they slept. Polysomnography (PSG, including EEG, electrooculogram, and electromyogram) and ECG were performed, and BF, respiratory activity, and oxygen saturation (SpO2) were measured. The PSG was performed using a PSG-1100 (Nihon Kohden, Tokyo, Japan) according to the AASM standards, and the sampling rate was set at 250 Hz. For the measurement of BF, customized sensors (KYOCERA, Tokyo, Japan) pre-calibrated to a “medical blood flow meter” (https://jpn.pioneer/ja/mhbd/bloodflow/) with a sampling rate of 39.0625 Hz were placed on the right index finger and the right ear concha. The measurement of BF was based on the LDF method. LDF provides a continuous estimate of skin BF restricted to the skin microcirculation and can be performed in different areas and on different surfaces.[21] The customized sensors used in this study can simultaneously estimate heartbeat activity (and the derived PRV) as well as BF, thus providing more information than photoplethysmography devices. Cellphones were used to store the BF data. The data were then exported to a computer for further analysis (Figure 1).

**Figure 1.**
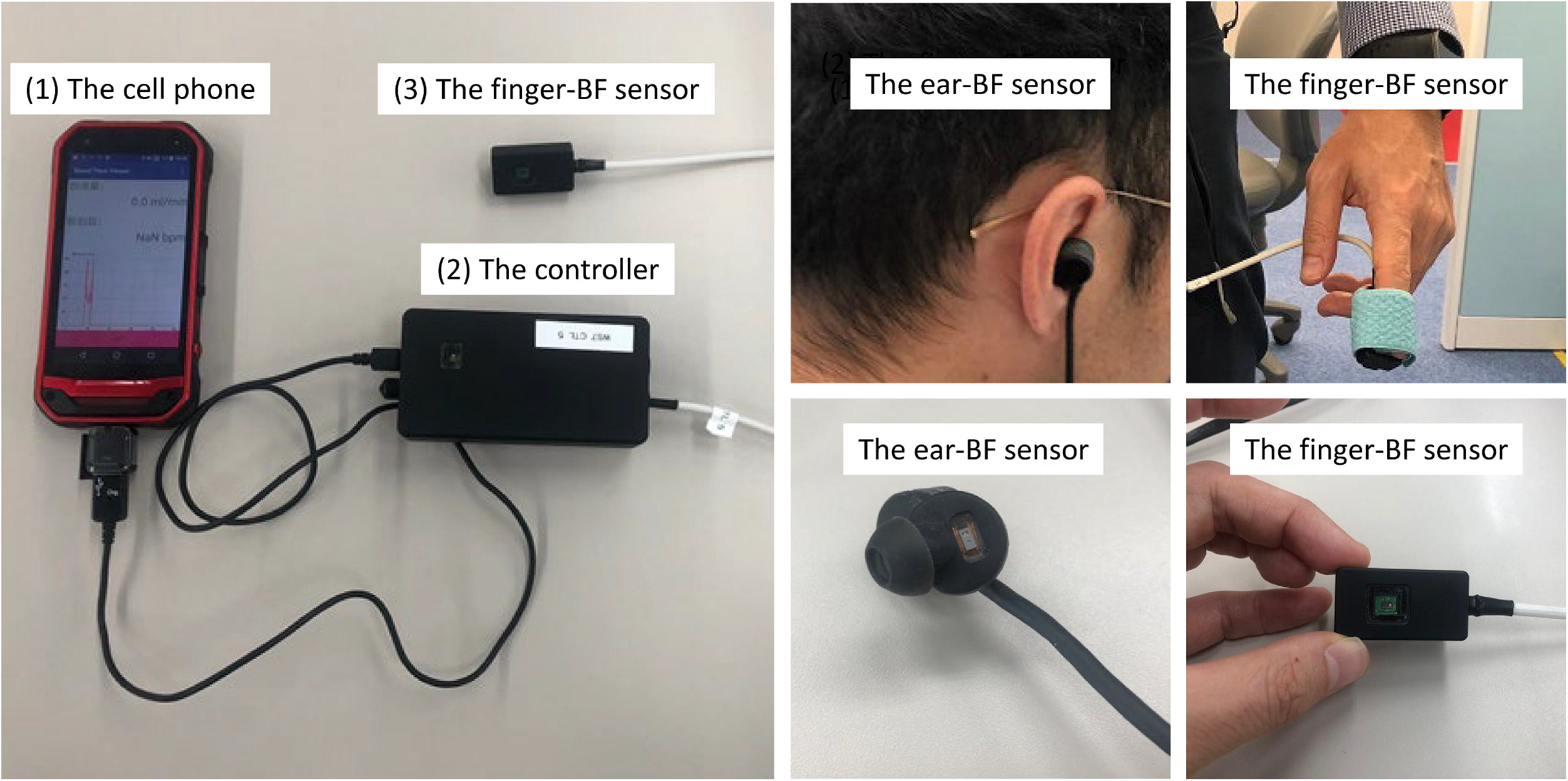
The devices used to measure finger- and ear-BF. BF, blood flow.

Respiratory activity was measured using a thermistor airflow (AF) sensor, a nasal pressure cannula, and a chest/abdomen band sensor compatible with the PSG-1100. SpO2 was also measured using a sensor compatible with the PSG-1100. In addition, we validated the accuracy of actigraphy (MotionWatch 8, CamNtech, Fenstanton, UK) compared to PSG (published elsewhere). The participants were asked to complete questionnaires about their sleep habits before sleeping. After sleeping, they completed a questionnaire about the experience of wearing the ear-BF sensor. Only PSG, ECG, BF, and AF data were used in this study. The data were analyzed using MATLAB (The Mathworks Inc., Natick, MA, USA), R (R Foundation for Statistical Computing, Vienna, Austria), and JASP (0.12.2; JASP Team, Amsterdam, Netherlands).

### 2.3 Data acquisition and preprocessing

All participants who passed the screening stage were required to sleep for eight hours in a sound-proof chamber between 20:00 to 8:00. PSG, finger-BF, ear-BF, AF, and other physiological activities were measured simultaneously from the same start time calibrated across devices. Thereafter, the data points were aligned between these four signals according to the start time and sampling rate. However, considering the availability of the data measured from the finger and ear sites, some participant data were excluded from further analysis. The exclusion criterion was less than two hours of BF data recorded during the night. After exclusion of some participants based on this criterion, the finger-BF data of 38 participants (10 women; mean age, 22.6 ± 3.7 years), ear-BF data of 42 participants (13 women; mean age, 22.6 ± 3.5 years), and the PSG, ECG, and AF data of 45 participants remained.

PSG data recorded during the 8-hour sleep after lights out were analyzed at 30-s epoch by 30-s epoch (in total, 960 epochs for each participant) by a registered polysomnographic technologist (Y. S.) for the sleep stages Wk, N1, N2, N3, and REM, according to the AASM criteria.[1] Finger- and ear-BF data and ECG data were analyzed based on EEG-defined sleep stages. However, considering the data length, most HRV indices require epochs longer than 1 min for the estimation of heart rate.[4] Therefore, we analyzed data based on 90-s epochs (in total, 320 epochs for each participant); that is, one 90-s epoch contained three consecutive 30-s epochs within the same sleep stage. Three consecutive epochs in the same sleep stage can reflect a relatively stable sleep stage, thus ensuring a stable relationship between the ANS and CNS in the same sleep stage. However, in the case of the three non-consecutive epochs, the 90-s epochs were set to be NaNs (not a number).

Before calculating the HRV/PRV indices, 90s-epochs of BF/ECG from each participant were preprocessed using a customized program in MATLAB. Data were first detrended and filtered using a onedimensional median filter to remove the spike noise that occurred during recording. The epochs with the largest amplitude beyond the three standard deviations (SDs) of the median value of the highest amplitudes were excluded to remove the epochs with significant artifacts. In addition, epochs with peaks of the heartbeat/pulse wave of BF/ECG (filtered using the default band-pass filter of 0.5–2 Hz embedded in the FieldTrip toolbox)[22] beyond the three SDs of the median value of all the peaks were also excluded to ensure the quality of the beat signal is optimal. The epochs were then visually inspected by the first author (Z. F.) to confirm their quality. The usability of the data after the preprocessing is shown in Table S1.

To derive the inter-beat intervals (IBIs) from the BF/ECG signals, peak detection was conducted on the waveforms of both ECG and BF signals using the “findpeaks” function in MATLAB. A pre-detection was first conducted to determine the parameter “MinPeakDistance” in “findpeaks”. ECG/BF signals were filtered using the default band-pass filter embedded in the FieldTrip toolbox with a band-pass frequency of 0.5–1.5 Hz. Peak detection was then performed using the default setting of “findpeaks” to determine the peaks and their locations (timings) from the filtered ECG/BF signals. For both BF and ECG signals, “MinPeakDistance” was set to 0.6*mean IBIs (differences in peak timings). For the formal peakdetection, BF signals were first filtered using the default high-pass filter embedded in the FieldTrip toolbox with a high-pass frequency of 0.5 Hz. Thereafter, the signals were then filtered using the “designfilt” function in MATLAB and a twelve order Butterworth low-pass filter with a cutoff frequency of 2.9297 Hz (0.15*sampling rate/2). We then used “findpeaks” to detect the peaks and their locations from the filtered BF signals, with “MinPeakDistance” determined using pre-detection, and “MinPeakHeight,” another parameter, set to zero. ECG signals were decomposed down to level 5 using the default ‘sym4’ wavelet by “modwt” function in MATLAB and then reconstructed by the “imodwt” function in MATLAB using only the wavelet coefficients at scales 4 and 5 (https://ww2.mathworks.cn/help/wavelet/ug/r-wave-detection-in-the-ecg.html). This was performed to enhance the R peaks in the ECG waveform. The reconstructed ECG signals were then subjected to peak detection using “findpeaks” with “MinPeakDistance” determined using pre-detection and “MinPeakHeight” set to 0.15*maximum amplitude of the reconstructed signals. These parameter settings were optimal for peak detection in this study, as confirmed by visual inspection of the detection results by the first author (Z. F.). Epochs with wrong peak detections of more than 10% (average of three wrong detections within one 30-s epoch) were excluded manually. Finally, the ectopic IBIs, which varied by more than 20% from the previous one (typically adopted in clinical practice and in human and animal research for HRV analysis),[23–25] were replaced with the mean value of the four neighboring IBIs centered on the ectopic one. However, the first two were replaced with the mean value of the four subsequent IBIs, whereas the last two were replaced with the mean value of the four previous IBIs, a method similar to that used in a previous study.[25]

We then calculated time-domain, frequency-domain, and non-linear HRV/PRV metrics related to ANS activity.[4] We selected several commonly used parameters in all three domains (Table S2). In the time domain, the SD of all the normal-to-normal (NN) intervals (i.e., typical IBIs resulting from sinus node depolarization,[18] which are free from artifacts) (SDNN), the root mean square of successive differences between adjacent NN intervals (RMSSD), and percentage of pairs of adjacent NN intervals differing by more than 50 ms (pNN50) are indices often chosen to reflect ANS activity.[4,18] The latter two indices measure cardiac vagal modulation.[3,4] In the frequency domain, generally within the frequency of 0.4 Hz, the HRV spectral power in the very low frequency range (< 0.04 Hz) is less likely to be interpreted. The HRV spectral power in the low frequency range (LF, 0.04 Hz–0.15 Hz) is often reported to reflect sympathetic activity; however, this conclusion is controversial.[3] In contrast, the high frequency range (HF, 0.15–0.4) reflects vagal (parasympathetic) activity.[3,4,18] LF and HF are usually normalized by dividing them by the total power (LF + HF). The normalized LF and HF (LFn and HFn) and the LF/HF ratio also reflect ANS activity [1]. There are also non-linear measures for the assessment of ANS activity using HRV and without assuming linearity and stationarity of the IBI time series. Such measures include detrended fluctuation analysis (DFA), which is a widely used for detecting short- and long-range correlations in non-stationary time series[4,26,27] and entropy measures, such as approximate entropy (ApEn),[4,28,29] which measures the predictability of fluctuations in the time series. These measures may assess non-linear HRV patterns related to more complex functioning, such as sleep. All indices, except for DFA1, DFA2, and ApEn, were calculated using a customized MATLAB script according to their definition. For DFA1, DFA2, and ApEn with more complex calculations, the scripts from the references[30,31] were used.

The power spectra of BF, PRV and HRV were analyzed using the “plomb” function in MATLAB. Since the data points of the HRV and PRV time series were sampled at different times, the frequency coordinates were different among epochs when calculating the power spectra. This made it impossible to average the power spectra across epochs in the same sleep stage or to compare the power spectra across sleep stages. Thus, interpolation was conducted after using “plomb” to unify the frequency coordinates. However, interpolation was not conducted on the BF power spectra. The power of a particular band within 0.2–0.3 Hz (as well as LFn, HFn, and LF/HF of BF) (Figure S1) was extracted, normalized by being divided by the HF, and compared. This frequency band is within the HF band (0.15–0.4 Hz), in which HRV power is modulated by sleep stages.[13,14] There are three reasons for focusing on this narrower band. First, we noted (without a hypothesis beforehand) that the group results of the BF, PRV and HRV spectra showed this peak frequency band. Second, previous studies have shown that stimulation around 0.25 Hz promotes sleep.[32–34] Third, the effects of respiration around 0.2–0.3 Hz on cardiovascular activity has been investigated in previous studies.[7,35]

To provide more information on the dynamics of BF, time-domain parameters, including the mean, SD, and the coefficient of variance (CV) of BF in different sleep stages, were calculated using the raw unfiltered BF data, but with epochs corresponding to PRV data after preprocessing.

For each index calculated for all participants, values larger than three SDs of the group means were set as missing values. Participants with missing values in either of the five stages were excluded from the group analysis. Thus, the group analysis for each dependent variable may have a different set of participants.

### 2.4 Statistical analysis

We investigated the changes in each HRV or (finger- or ear-) PRV parameter across the sleep stages separately for each recording site (heart, finger, or ear). The Jarque-Bera test[36] with a significance level of 0.01 was used to test the normality of the distribution of each index across individuals. For HRV/PRV indices that met the normality, we performed parametric analysis using one-way (stages: Wk, N1, N2, N3, and REM) frequentist repeated measures analysis of variance (F-RMANOVA) with Greenhouse–Geisser (GG) correction for the violation of sphericity and its Bayesian version (B-RMANOVA) for significance testing. For those that did not meet normality, we conducted non-parametric analysis using the Friedman’s test with the Conover’s post-hoc test for pairwise comparisons. The Bayesian Wilcoxon signed-rank test was conducted as a non-parametric version of the Bayesian pairwise comparisons. To thoroughly provide information about the differences in HRV/PRV indices across sleep stages, a post-hoc test was conducted for all possible pairs of sleep stages. It was corrected for multiple (ten in total) comparisons using the Bonferroni method. Statistical significance was set at *p* < 0.05. However, for the Bayesian analysis approach, the Bayes factors were derived and interpreted using a classification scheme.[37–39] The advantage of using the Bayes factor is that it shows the amount of evidence for the null hypothesis (H0) or the alternative hypothesis (H1), or insufficient evidence for either hypothesis.[40] Therefore, even if F-RMANOVA does not yield significant results, which means that H0 can neither be rejected nor accepted, the Bayes factor can still provide the amount of evidence for H1 against H0. For example, the Bayes factor *B*_10_ shows the level of the possibility of H1 against H0, and its value is classified into different categories of evidence (Table S3, adapted from[38,39]). Further, if there is more than one alternative hypothesis or model (both terms are used interchangeably in this study) other than H1, say H2 (for an example of more than one hypothesis, see[37]), against H0, Bayes factors that compare H1 and H2 can also indicate which model (H1 or H2) is more supported. When H1 and H2 correspond to different combinations of the factors, the winning of either hypothesis can indicate which of the two combinations of factors is preferred.

We investigated the similarities or differences in the patterns of PRV/HRV indices recorded in the different sleep stages and at the different sites (finger, ear, and heart). For the indices that met the normality, we performed parametric analysis using a two-way (stage: Wk, N1, N2, N3, and REM; site: heart, finger, ear) F-RMANOVA and its B-RMANOVA for significance testing, with GG correction for the violation of sphericity. For those that did not meet normality, the data were transformed using the boxcox method into normally distributed data and subjected to parametric analysis. The parametric analysis provided more detailed information, such as the effect size. However, for those that were difficult to transform, we conducted non-parametric two-way analysis using R with the software package nparLD.[41,42] We focused on the interaction between stage and site because it reveals the similarities and differences in patterns recorded in the different sleep stages and at the different sites (heart, finger, and ear).

For the BF parameters, we only investigated the changes in each parameter across the sleep stages separately for each recording site (finger or ear) using methods similar to those used for PRV parameters.

## 3 Results

Although data of 45 participants (13 women; mean age, 22.5 ± 3.4 years) were obtained, the finger-BF data of 38 participants (10 women; mean age, 22.6 ± 3.7 years), ear-BF data of 42 participants (13 women; mean age, 22.6 ± 3.5 years), and the PSG, ECG, and AF data of 45 participants remained after preprocessing. However, for each index investigated as follows, values larger than three SDs of the group means were set as missing values. Participants with missing values in either of the five stages were excluded from the group analysis. Thus, the group analysis for each dependent variable may have a different set of participants. For example, for heat-IBIs, data of only forty participants (12 women) remained for group analysis. The details are shown in the following results.

### 3.1 Comparison of changes in finger- and ear-PRV and HRV indices across different sleep stages

Figure 2 shows the mean heart-, finger-, and ear-IBIs across the different sleep stages. The results showed the main effects of stage on mean heart-IBIs (*N* = 40, 12 women; mean age, 22.6 ± 3.5 years; *F*(4, 156) = 46.48; *p* < 0.001; 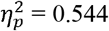, GG-corrected for non-sphericity), mean finger-IBIs (*N* = 27, 8 women; mean age, 22.0 ± 1.2 years; *F*(4, 104) = 22.15; *p* < 0.001; 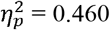, GG-corrected), and mean ear-IBIs (*N* = 23, 6 women; mean age, 22.7 ± 4.6 years; *F*(4, 88) = 29.67; *p* < 0.001; 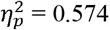). These results showed that the mean IBI was different across sleep stages, providing information for differentiating all pairs of different sleep stages, except N1 and REM, for all three recording sites (see Table S4). The mean IBI was higher (i.e., heart rate was lower) in NREM sleep (specifically, N2 and N3) than in REM sleep, and in sleep than in Wk.

**Figure 2.**
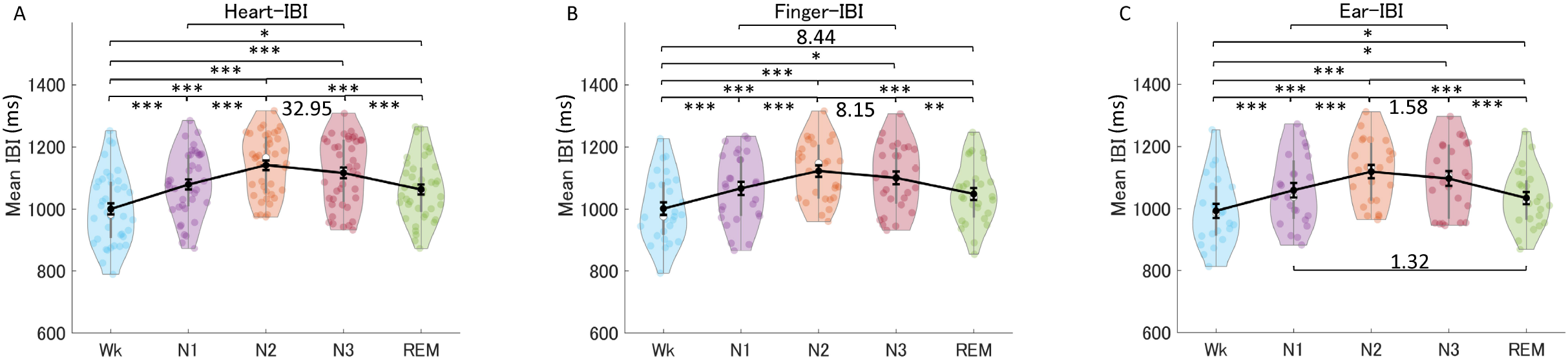
Mean IBIs across the different sleep stages for the heart (A), the finger (B), and the ear (C). The violin plot with dots shows the distribution of the individual data points. The line chart with error bars shows the group mean and the ± 1 standard error of the mean. \**p* < 0.05; \*\**p* < 0.01; \*\*\**p* < 0.001. The numerical values are the Bayes factors. Only the values for nonsignificant comparisons were provided here for complementary information (for other information, see Table S4). The values show anecdotal (1–3), moderate (3–10), or strong (>10) evidence against the H0 of no difference between pairs of sleep stages. Bayes factors < 1 are not listed. IBI, inter-beat interval.

The results of comparisons of the patterns of the three recording sites across five different sleep stages showed the main effects of stage (*N* = 14, 3 women; mean age, 21.6 ± 1.2 years; *F*(4, 52) = 29.50; *p* < 0.001; 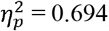) and site (*F*(2, 26) = 7.83; *p* = 0.007; 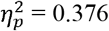, GG-corrected). There was a marginally significant interaction between the two factors (*F*(8, 104) = 2.30; *p* = 0.090; 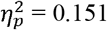, GG-corrected). The Bayes factors for model comparison (Table S5) showed that compared with the null model (*B*_10_ > 100), the model with the stage factor only was the best and outperformed the other models. The nonsignificant interaction between stage and site, and the best model without the “stage×site” factor, revealed similar patterns across the sleep stages for the mean heart-IBI, mean finger-IBI, and mean ear-IBI.

#### 3.1.1 Time-domain indices

Figure 3 shows the HRV, finger-PRV, and ear-PRV indices in the time domain, including SDNN, RMSSD, and pNN50 (Table S4). For the SDNN of each recording site, the results showed the main effects of stage for HRV (*N* = 40, 12 women; mean age, 22.7 ± 3.6 years; *F*(4, 156) = 15.45; *p* < 0.001; 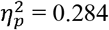, GG-corrected), finger-PRV (*N* = 27, 8 women; mean age, 22.1 ± 1.4 years; *F*(4, 104) = 9.31; *p* < 0.001; 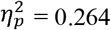, GG-corrected), and ear-PRV (*N* = 23, 6 women; mean age, 22.7 ± 4.6 years; *F*(4, 88) = 2.64; *p* = 0.039; 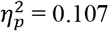). The results of comparisons of the patterns of the three recording sites across five different sleep stages showed the main effects of stage (*N* = 14, 3 women; mean age, 21.6 ± 1.2 years; *F*(4, 52) = 6.37; *p* < 0.001; 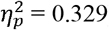) and site (*F*(2, 26) = 96.14; *p* < 0.001; 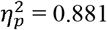, GG-corrected). There was no significant interaction between the two factors (*F*(8, 104) = 1.82; *p* = 0.148; 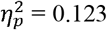, GG-corrected). The Bayes factors for model comparison (Table S5) showed that compared with the null model (*B*_10_ > 100), the model with the factors “stage+site+stage×site” was the best and far outperformed other models. Although the interaction between stage and site was not significant, the “stage×site” component in the best model suggested there might be a trend of different, but non-significant, SDNN patterns across the heart, finger, and ear sites.

**Figure 3.**
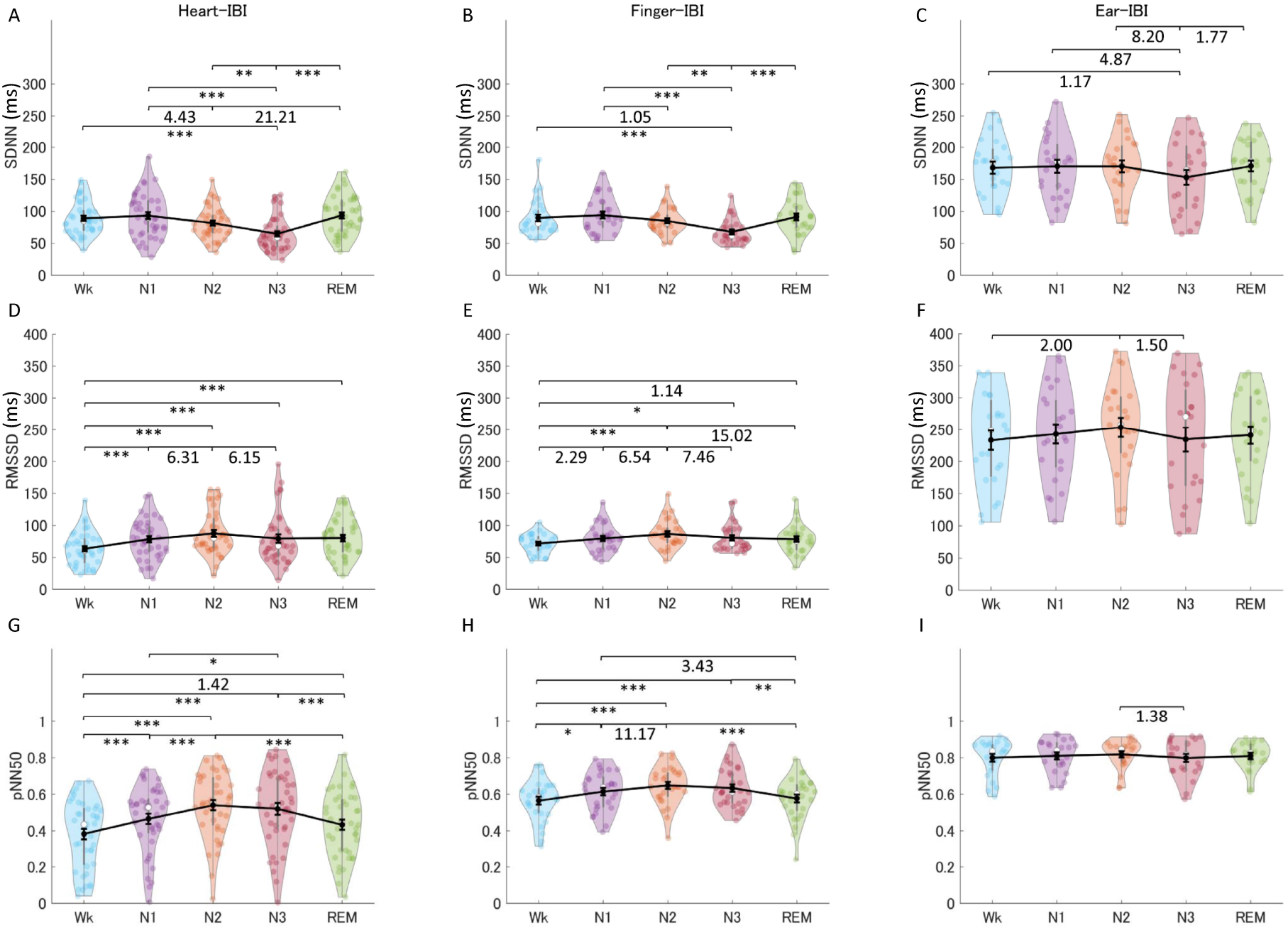
The indices of HRV (A, D, G), finger-PRV (B, E, H), and ear-PRV (C, F, I) in the time domain, including SDNN (A-C), RMSSD (D-F), and pNN50 (G-I), across the different sleep stages. The violin plot with dots shows the distribution of the individual data points. The line chart with error bars shows the group mean and the ± 1 standard error of the mean. *: *p* < 0.05; **: *p* < 0.01; \*\*\**p* < 0.001. The numerical values are the Bayes factors. Only those for nonsignificant comparisons were provided here for complementary information (for other information, see Table S4). The values show anecdotal (1–3), moderate (3–10), or strong (> 10) evidence against the H0 of no difference between pairs of sleep stages. Bayes factors < 1 are not listed. HRV, heart rate variability; PRV, pulse rate variability; SDNN, standard deviation of all the normal-to-normal intervals; RMSSD, root mean square of successive differences between adjacent normal-to-normal intervals; pNN50, percentage of pairs of adjacent normal-to-normal intervals differing by more than 50 ms; IBI, inter-beat interval.

For RMSSD, the results showed the main effects of stage for HRV (*N* = 38, 11 women; mean age, 22.7 ± 3.7 years; *F*(4, 148) = 10.00; *p* < 0.001; 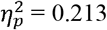, GG-corrected), finger-PRV (*N* = 27, 8 women; mean age, 22.1 ± 1.4 years; *F*(4, 104) = 6.60; *p* < 0.001; 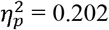, GG-corrected), and ear-PRV (*N* = 23, 6 women; mean age, 22.7 ± 4.6 years; *F*(4, 88) = 1.33; *p* = 0.273; 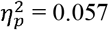, GG-corrected). The results of comparisons of the patterns of the three recording sites across five different stages showed the main effects of stage (*N* = 14, 3 women; mean age, 21.6 ± 1.2 years; *F*(4, 52) = 3.16; *p* = 0.021; 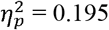) and site (*F*(2, 26) = 134.04; *p* < 0.001; 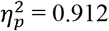, GG-corrected). There was no significant interaction between the two factors (*F*(8, 104) = 0.38; *p* = 0.755; 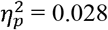; GG-corrected). The Bayes factors for model comparison (Table S5) showed that compared with the null model (*B*_10_ > 100), the model with site was the best and outperformed the other models. The non-significant interaction between stage and site, as well as the best model without the factor “stage×site”, revealed the same patterns across sleep stages for HRV, finger-PRV, and ear-PRV.

For pNN50, the results showed the main effects of stage for HRV (*N* = 41, 12 women; mean age, 22.7 ± 3.5 years; *F*(4, 160) = 24.69; *p* < 0.001; 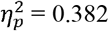, GG-corrected), finger-PRV (*N* = 28, 8 women; mean age, 22.1 ± 1.4 years; *F*(4, 108) = 10.28; *p* < 0.001; 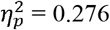, GG-corrected), and ear-PRV (*N* = 22, 6 women; mean age, 22.7 ± 4.7 years; *F*(4, 84) = 0.80; *p* = 0.528; 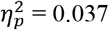). The results of comparisons of the patterns of the three recording sites across five different stages showed the main effects of stage (*N* = 14, 3 women; mean age, 21.6 ± 1.2 years; *F*(4, 52) = 7.39; *p* < 0.001; 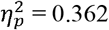) and site (*F*(2, 26) = 131.81,*p* < 0.001, 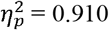), and the interaction between the two factors (*F*(8, 104) = 7.23, *p* = 0.001, 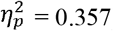). The Bayes factors for model comparison (Table S5) showed that compared with the null model (*B*_10_ > 100), the model with “stage+site+stage×site” was the best and far outperformed the other models. The significant interaction between stage and site, and the best model with the interaction factor “stage×site”, revealed different patterns for HRV, finger-PRV, and ear-PRV across sleep stages. The results of post-hoc tests for the comparison of pNN50 between HRV and finger-HRV showed the main effects of stage (*N* = 28, 8 women; mean age, 22.1 ± 1.4 years; *F*(4, 108) = 14.14; *p* < 0.001; 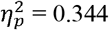, GG-corrected) and site (*F*(1, 27) = 74.74,*p* < 0.001, 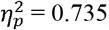), and the interaction between the two factors (*F*(4, 108) = 7.55, *p* = 0.001, 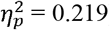). The Bayes factors for model comparison showed that compared with the null model (*B*_10_ > 100), the model with “stage+site” was the best and outperformed the other models. Although the interaction between stage and site was significant, the best model did not have the interaction factor “stage×site”. This revealed that although the patterns of HRV and finger-PRV across sleep stages are different, the difference might be relatively trivial compared with the main effects of stage and site. The results of post-hoc tests for the comparison of pNN50 between HRV and ear-HRV showed the main effects of stage (*N* = 22, 6 women; mean age, 22.7 ± 4.7 years; *F*(4, 84) = 13.56; *p* < 0.001; 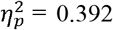, GG-corrected) and site (*F*(1, 21) = 198.57, *p* < 0.001, 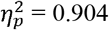), and the interaction between the two factors (*F*(4, 84) = 17.89, *p* = 0.001, 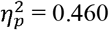, GG-corrected). The Bayes factors for model comparison showed that compared with the null model (*B*_10_ > 100), the model with “stage+site+stage×site” was the best and far outperformed the other models. The significant interaction between stage and site, and the best model with the interaction factor “stage×site”, revealed that the HRV and ear-PRV patterns across sleep stages were significantly different. These results implied that although there was a difference in pNN50 between HRV and PRV, the pattern of finger-pNN50 across the sleep stages was more similar to that of heart-pNN50 than that of ear-pNN50 (Table S4).

Analysis of the results of the PRV/HRV indices in the time domain showed that the finger indices have very similar patterns to those of the heart. Specifically, the SDNN for HRV and finger-PRV were lower in NREM (N3) sleep than in REM sleep and Wk. For NREM sleep stages, SDNN was lower in N3 than in N2 and N1, but lower in N2 than in N1 (weak evidence). Ear-SDNN showed similar results, but with weaker evidence. The RMSSD for HRV and finger-PRV were higher in NREM than in Wk. For NREM sleep, moderate evidence observed from Bayes factors showed that heart- and finger-RMSSD were higher in N2 than in N1 and N3. Ear-RMSSD also showed anecdotal evidence to be highest in N2. Similarly, the pNN50 for HRV and finger-PRV were higher during NREM sleep than during Wk and REM sleep. For NREM sleep, heart- and finger-RMSSD were higher in N2 than in N1. Therefore, the sensitivity of PRV may be lower than that of HRV, as reflected by the less significant differences and small Bayes factors between the different sleep stages.

#### 3.1.2 Frequency-domain indices

Figure 4 shows the indices of HRV, finger-PRV, and ear-PRV in the frequency domain, including LFn, HFn, and LF/HF (Table S4). For the LFn of each recording site, the results showed the main effects of stage for HRV (*N* = 40, 12 women; mean age, 22.7 ± 3.5 years; *F*(4, 160) = 73.52; *p* < 0.001; 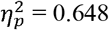, GG-corrected), finger-PRV (*N* = 28, 8 women; mean age, 22.1 ± 1.4 years; *F*(4, 108) = 27.24; *p* < 0.001; 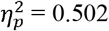, GG-corrected), and ear-PRV (*N* = 22, 5 women; mean age, 22.6 ± 4.7 years; *F*(4, 84) = 7.83; *p* < 0.001; 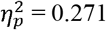). The results of comparisons of the patterns of the three recording sites across five different stages showed the main effects of stage (*N* = 13, 2 women; mean age, 21.5 ± 1.2 years; *F*(4, 48) = 25.41; *p* < 0.001; 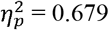, GG-corrected) and site (*F*(2, 24) = 88.66, *p* < 0.001, 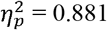), and the interaction between the two factors (*F*(8, 96) = 11.40; *p* < 0.001; 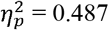, GG-corrected). The Bayes factors for model comparison (Table S5) showed that compared with the null model (*B*_10_ > 100), the model with “stage+site+stage×site” was the best and far outperformed the other models. The significant interaction between stage and site, and the best model with the interaction factor “stage×site”, revealed different patterns for HRV, finger-PRV, and ear-PRV across sleep stages. The results of post-hoc tests for the comparison of LFn between HRV and finger-HRV showed the main effects of stage (*N* = 28, 8 women; mean age, 22.1 ± 1.4 years; *F*(4, 108) = 47.93; *p* < 0.001; 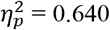, GG-corrected) and site (*F*(1, 27) = 8.81, *p* = 0.006, 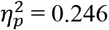), and the interaction between the two factors (*F*(4, 108) = 5.00, *p* = 0.004, 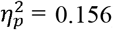). The Bayes factors for model comparison showed that compared with the null model (*B*_10_ > 100), the model with “stage+site” was the best and outperformed other models. Although the interaction between stage and site was significant, the best model did not have the interaction factor “stage×site”. Furthermore, although the HRV and finger-PRV patterns were different across sleep stages, the difference might be relatively trivial compared with the main effects of stage and site. The results of post-hoc tests for the comparison of LFn between HRV and ear-HRV showed the main effects of stage (*N* = 22, 5 women; mean age, 22.6 ± 4.7 years; *F*(4, 84) = 30.21; *p* < 0.001; 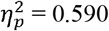, GG-corrected) and site (*F*(1, 21) = 149.78, *p* < 0.001, 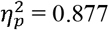), and the interaction between the two factors (*F*(4, 84) = 24.48, *p* < 0.001, 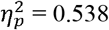, GG-corrected). The Bayes factors for model comparison showed that compared with the null model (*B*_10_ > 100), the model with “stage+site+stage×site” was the best and far outperformed the other models. The significant interaction between stage and site, and the best model with the interaction factor “stage×site”, revealed that the HRV and ear-PRV patterns across sleep stages were significantly different. These results implied that although there was a difference in LFn between HRV and PRV, the pattern of finger-LFn across the sleep stages was more similar to that of the heart-LFn than that of the ear-LFn.

**Figure 4.**
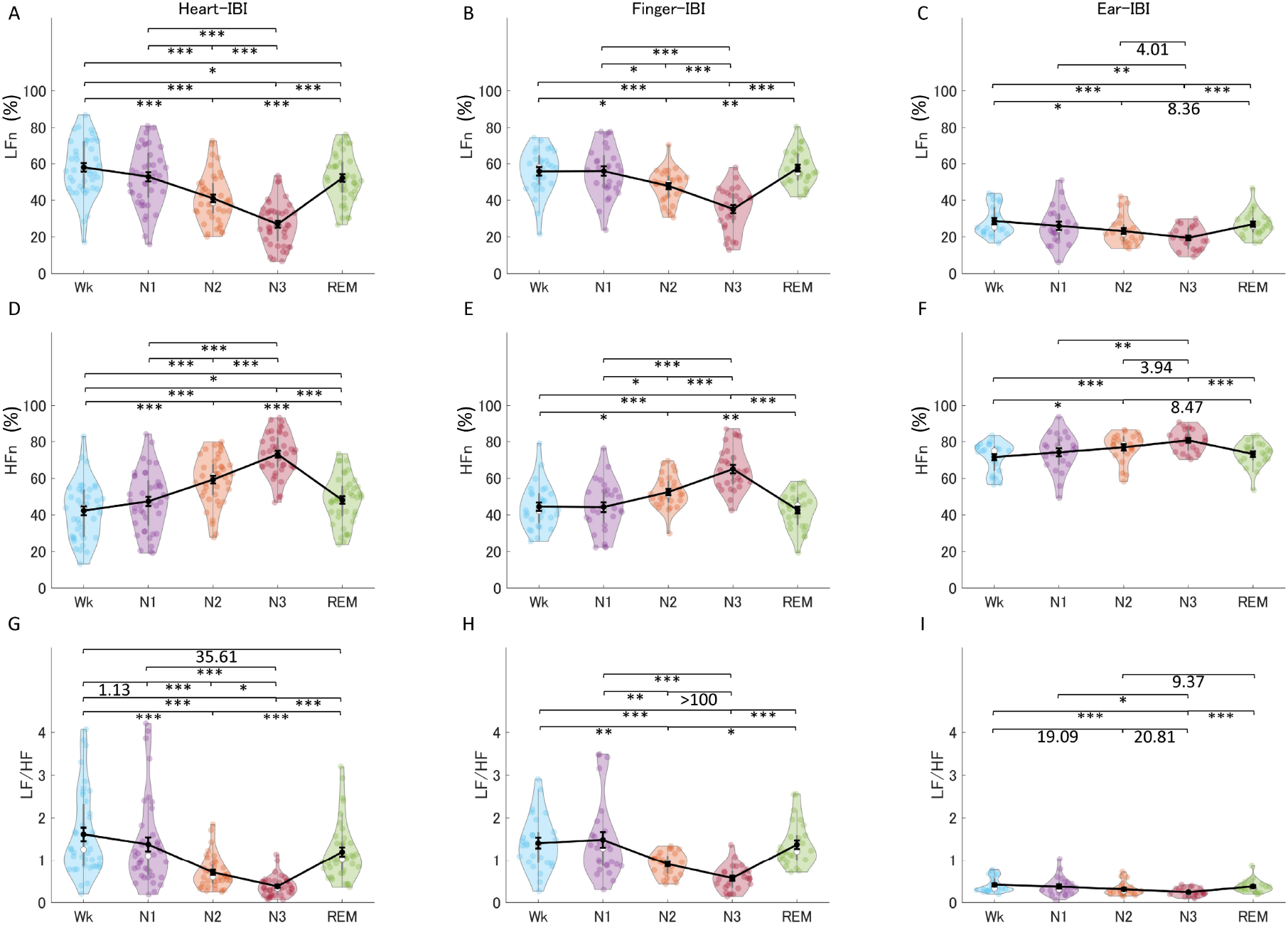
The indices of HRV (A, D, G), finger-PRV (B, E. H), and ear-PRV (C, F, I) in the frequency domain, including LFn (A-C), HFn (D-F), and LF/HF (G-I), across the different sleep stages. The violin plot with dots shows the distribution of the individual data points. The line chart with error bars shows the group mean and the ± 1 standard error of the mean. \**p* < 0.05; \*\**p* < 0.01; \*\*\**p* < 0.001. The numerical values are the Bayes factors. Only those for nonsignificant comparisons were provided here for complementary information (for other information, see Table S4). The values show anecdotal (1–3), moderate (3–10), strong (> 10), or extreme evidence (> 100) against the H0 of no difference between pairs of sleep stages. Bayes factors < 1 are not listed. HRV, heart rate variability; PRV, pulse rate variability; LF, low frequency power; HF, high frequency power; LFn, normalized low frequency power; HFn, normalized high frequency power; IBI, inter-beat interval.

Regarding the HFn of each recording site, the results showed the main effects of stage for HRV (*N* = 41, 12 women; mean age, 22.7 ± 3.5 years; *F*(4, 160) = 73.17;*p* < 0.001; 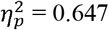, GG-corrected), finger-PRV (*N* = 28, 8 women; mean age, 22.1 ± 1.4 years; *F*(4, 108) = 27.11; *p* < 0.001; 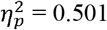, GG-corrected), and ear-PRV (*N* = 22; 5 women; mean age, 22.6 ± 4.7 years; *F*(4, 84) = 7.80; *p* < 0.001; 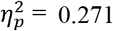). The results of comparisons of the patterns of the three recording sites across five different stages showed the main effects of stage (*N* = 13, 2 women; mean age, 21.5 ± 1.2 years; *F*(4, 48) = 25.37; *p* < 0.001; 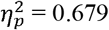, GG-corrected) and site (*F*(2, 24) = 89.04, *p* < 0.001, 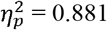), and the interaction between the two factors (*F*(8, 96) = 11.35, *p* < 0.001, 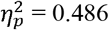, GG-corrected). The Bayes factors for model comparison (Table S5) showed that compared with the null model (*B*_10_ > 100), the model with “stage+site+stage×site” was the best and far outperformed the other models. The significant interaction between stage and site, and the best model with the interaction factor “stage×site”, revealed different patterns of HRV, finger-PRV, and ear-PRV across sleep stages. The results of post-hoc tests for the comparison of HFn between HRV and finger-HRV showed the main effects of stage (*N* = 28, 8 women; mean age, 22.1 ± 1.4 years; *F*(4, 108) = 47.64; *p* < 0.001; 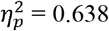, GG-corrected) and site (*F*(1, 27) = 8.77, *p* = 0.006, 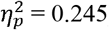), and the interaction between the two factors (*F*(4, 108) = 5.00, *p* = 0.004, 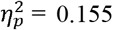, GG-corrected). The Bayes factors for model comparison showed that compared with the null model (*B*_10_ > 100), the model with “stage+site” was the best and outperformed other models. Although the interaction between stage and site was significant, the best model did not have the interaction factor “stage×site”. In addition, although the patterns for HRV and finger-PRV were different across sleep stages, the difference might be relatively trivial compared with the main effects of stage and site. The results of post-hoc tests for the comparison of HFn between HRV and ear-HRV showed the main effects of stage (*N* = 22, 5 women; mean age, 22.6 ± 4.7 years; *F*(4, 84) = 30.17; *p* < 0.001; 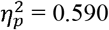, GG-corrected) and site (*F*(1, 21) = 150.75, *p* < 0.001, 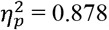), and the interaction between the two factors (*F*(4, 84) = 24.35, *p* < 0.001, 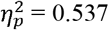, GG-corrected). The Bayes factors for model comparison showed that compared with the null model (*B*_10_ > 100), the model with “stage+site+stage×site” was the best and far outperformed the other models. The significant interaction between stage and site, and the best model with the interaction factor “stage×site”, revealed that the patterns for HRV and ear-PRV across sleep stages were significantly different. These results implied that although there was a difference in HFn between HRV and PRV regarding HFn, the pattern of finger-HFn across the sleep stages was more similar to that of heart-HFn than that of ear-HFn (Table S4).

Regarding the LF/HF of each recording site, the results showed the main effects of stage for HRV (*N* = 39, 12 women; mean age, 22.7 ± 3.6 years; *χ*^2^(4) = 104.31;*p* < 0.001), finger-PRV (*N* = 26, 7 women; mean age, 22.2 ± 1.5 years; *F*(4, 100) = 14.90; *p* < 0.001; 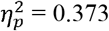, GG-corrected), and ear-PRV (*N* = 22, 5 women; mean age, 22.6 ± 4.7 years; *χ*^2^(4) = 30.98; *p* < 0.001). The LF/HF data were transformed using the boxcox method into normally distributed data and then subjected to parametric analysis because it violated the normality. The results of comparisons of the patterns across five different stages for the three recording sites showed the main effects of stage (*N* = 11, 11 woman; mean age, 21.5 ± 1.3 years; *F*(4, 40) = 20.61; *p* < 0.001; 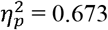, GG-corrected) and site (*F*(2, 20) = 65.70, *p* < 0.001, 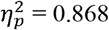), and the interaction between the two factors (*F*(8, 80) = 9.00, *p* < 0.001, 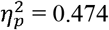, GG-corrected). The Bayes factors for model comparison (Table S5) showed that compared with the null model (*B*_10_ > 100), the model with “stage+site+stage×site” was the best and far outperformed the other models. The significant interaction between stage and site, and the best model with the interaction factor “stage×site”, revealed different patterns for HRV, finger-PRV, and ear-PRV across sleep stages. The results of post-hoc tests for the comparison of LF/HF between HRV and finger-HRV showed the main effects of stage (*N* = 26, 7 women; mean age, 22.2 ± 1.5 years; *F*(4, 100) = 45.10; *p* < 0.001; 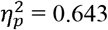, GG-corrected) and site (*F*(1, 25) = 8.83, *p* = 0.006, 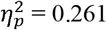), and the interaction between the two factors (*F*(4, 100) = 5.61, *p* = 0.003, 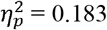, GG-corrected). The Bayes factors for model comparison showed that compared with the null model (B_10_ > 100), the model with “stage+site” was the best and outperformed other models. Although the interaction between stage and site was significant, the best model did not have the interaction factor “stage×site”. Furthermore, although the patterns for HRV and finger-PRV were different across sleep stages, the difference was relatively trivial compared to the main effects. The results of post-hoc tests for the comparison of LF/HF between HRV and ear-HRV showed the main effects of stage (*N* = 22, 5 women; mean age, 22.6 ± 4.7 years; *F*(4, 76) = 24.44; *p* < 0.001; 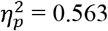, GG-corrected) and site (*F*(1, 19) = 123.10, *p* < 0.001, 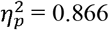), and the interaction between the two factors (*F*(4, 76) = 18.36, *p* < 0.001, 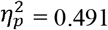). The Bayes factors for model comparison showed that compared with the null model (*B*_10_ > 100), the model with “stage+site+stage×site” was the best and far outperformed the other models. The significant interaction between stage and site, and the best model with the interaction factor “stage×site”, revealed that the patterns for HRV and ear-PRV were significantly different across sleep stages. These results imply that although LF/HF was different between HRV and PRV, the pattern of finger-LF/HF across the sleep stages was more similar to that of the heart-LF/HF than that of the ear-LF/HF.

Like the results of the PRV/HRV indices in the time domain, the finger indices in the frequency domain showed very similar patterns to those of the heart indices. The ear indices showed fewer significant comparisons between sleep stages. Specifically, LFn for HRV, finger-PRV, and ear-PRV were lower in NREM (N2 and N3) sleep, REM sleep, and Wk. For NREM sleep, LFn was lower in N3 than in N2 and N1, and lower in N2 than in N1 (except for ear-LFn). HFn showed the opposite results to those of LFn, whereas LF/HF showed similar results to those of LFn.

#### 3.1.3 Non-linear measurements

Figure 5 shows the non-linear indices of HRV, finger-PRV, and ear-PRV, including ApEn, DFA1, and DFA2 (Table S4). For the ApEn of each recording site, the results showed the main effects of stage for HRV (*N* = 39, 12 women; mean age, 22.6 ± 3.6 years; *F*(4, 142) = 45.35; *p* < 0.001; 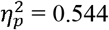, GG-corrected), finger-PRV (*N* = 26, 7 women; mean age, 22.0 ± 1.3 years; *χ*^2^(4) = 17.20; *p* = 0.002), and ear-PRV (*N* = 21, 5 women; mean age, 22.6 ± 4.8 years; *F*(4, 80) = 0.99; *p* = 0.400; 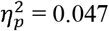, GG-corrected). ApEn data were transformed using the boxcox method into normally distributed data and then subjected to parametric analysis because it violated the normality. The results of the comparisons of the patterns of the three recording sites across five different stages showed the main effect of stage (*N* = 12, 2 women; mean age, 21.4 ± 1.1 years; *F*(4, 44) = 7.24; *p* < 0.001; 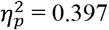), the marginally significant effect of site (*F*(2, 22) = 3.72; *p* = 0.062; 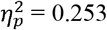, GG-corrected), and the marginally significant interaction between the two factors (*F*(8, 88) = 2.60; *p* = 0.057; 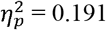, GG-corrected). The Bayes factors for model comparison (Table S5) showed that compared to the null model (*B*_10_ > 100), the model with “stage+site+stage×site” was the best and outperformed other models; however, the second-best model with “stage+site” was very close to the best model (*B* = 0.71, compared with the best model). The marginally significant interaction between stage and site, and the best model with the interaction factor “stage×site”, suggested a trend of different patterns of HRV, finger-PRV, and ear-PRV across sleep stages.

**Figure 5.**
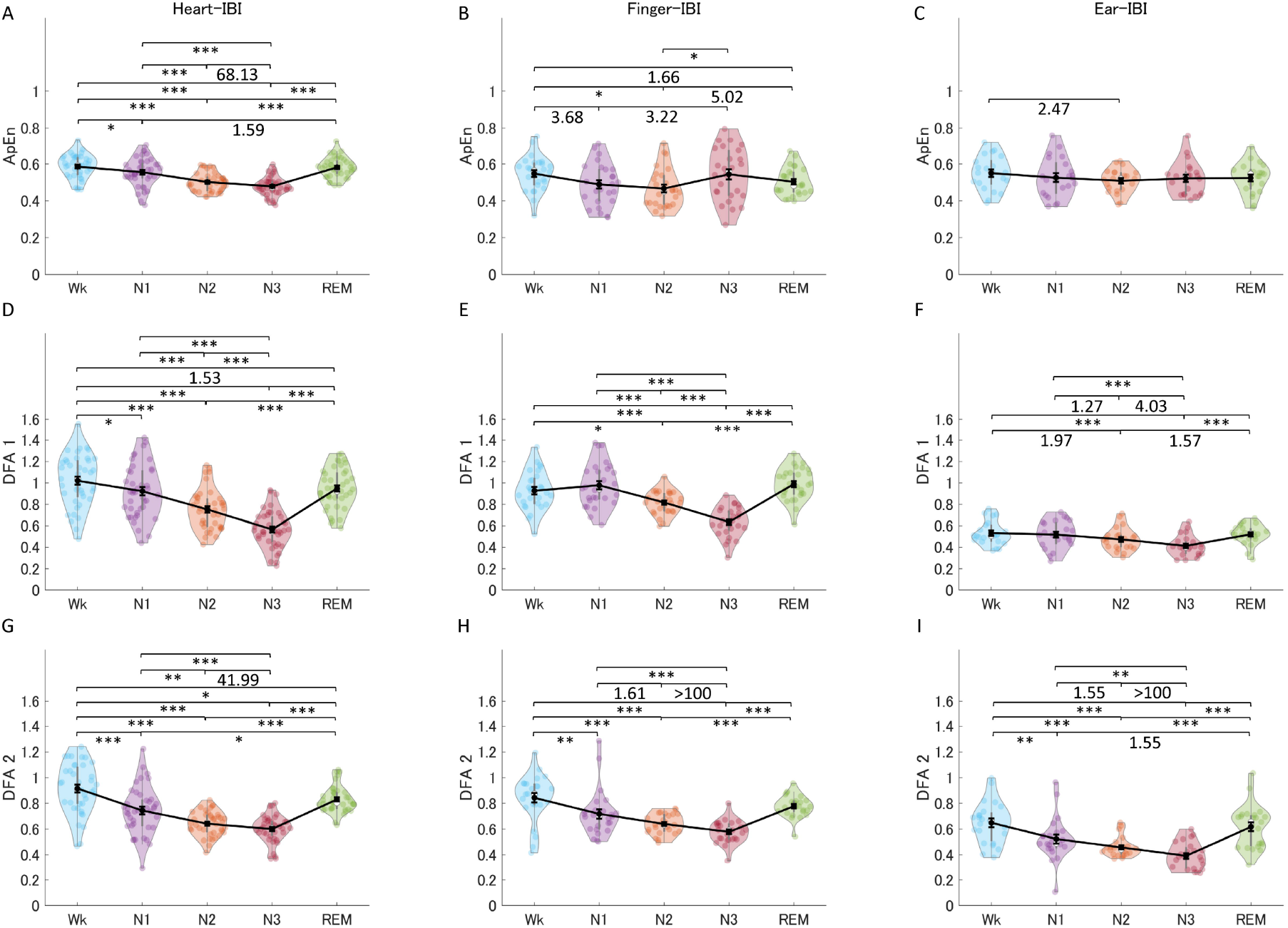
The indices of HRV (A, D, G), finger-PRV (B, E. H), and ear-PRV (C, F, I) in non-linear measurements, including ApEn (A-C), DFA1 (D-F), and DFA2 (G-I), across the different sleep stages. The violin plot with dots shows the distribution of the individual data points. The line chart with error bars shows the group mean and the ± 1 standard error of the mean. \**p* < 0.05; \*\**p* < 0.01; \*\*\**p* < 0.001. The numerical values are the Bayes factors. Only those for nonsignificant comparisons were provided here for complementary information (for other information, see Table S4). The values show anecdotal (1–3), moderate (3–10), strong (> 10), or extreme evidence (> 100) against the H0 of no difference between pairs of sleep stages. Bayes factors < 1 are not listed. HRV, heart rate variability; PRV, pulse rate variability; ApEn, approximate entropy; DFA, detrended fluctuation analysis; IBI, inter-beat interval.

For the DFA1 of each recording site, the results showed the main effects of stage for HRV (*N* = 41, 12 women; mean age, 22.7 ± 3.5 years; *F*(4, 160) = 69.95;*p* < 0.001; 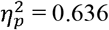, GG-corrected), finger-PRV (*N* = 28, 8 women; mean age, 22.1 ± 1.4 years; *F*(4, 108) = 35.64; *p* < 0.001; 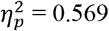, GG-corrected), and ear-PRV (*N* = 22, 5 women; mean age, 22.6 ± 4.7 years; *F*(4, 84) = 8.69; *p* < 0.001; 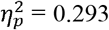). The results of comparisons of the patterns of the three recording sites across five different sleep stages showed the main effects of stage (*N* = 13, 2 women; mean age, 21.5 ± 1.2 years; *F*(4, 48) = 31.10; *p* < 0.001; 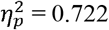) and site (*F*(2, 24) = 126.2, *p* < 0.001, 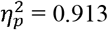), and the interaction between the two factors (*F*(8, 96) = 9.71, *p* < 0.001, 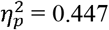). The Bayes factors for model comparison (see Table S5) showed that the model with “stage+site+stage×site” was the best compared with the null model (*B*_10_ > 100) and far outperformed the other models. The significant interaction of stage and site, and the best model with the interaction factor “stage×site”, revealed different patterns of HRV, finger-PRV, and ear-PRV across sleep stages. The results of post-hoc tests for the comparison of DFA1 between HRV and finger-HRV showed the main effects of stage (*N* = 28, 8 women; mean age, 22.1 ± 1.4 years; *F*(4, 108) = 54.87; *p* < 0.001; 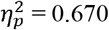, GG-corrected) and the interaction between stage and site (*F*(4, 108) = 4.60, *p* = 0.007, 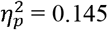, GG-corrected). No significant effect of site was noted (*F*(1, 27) = 1.08, *p* = 0.308, 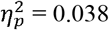). The Bayes factors for model comparison showed that compared with the null model (*B*_10_ > 100), the model with stage was the best and outperformed the other models. Although the interaction between stage and site was significant, the main effect of site was not significant. The best model did not have the interaction factor “stage×site”, which suggests that although the HRV and finger-PRV patterns across sleep stages were different, the difference was relatively trivial compared with the main effects of stage. The results of post-hoc tests for the comparison of DFA1 between HRV and ear-HRV showed the main effects of stage (*N* = 22, 5 women; mean age, 22.6 ± 4.7 years; *F*(4, 84) = 31.90; *p* < 0.001; 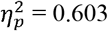, GG-corrected) and site (*F*(1, 21) = 203.351, *p* < 0.001, 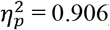), and the interaction between the two factors (*F*(4, 84) = 21.50; *p* < 0.001; 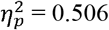, GG-corrected). The Bayes factors for model comparison showed that compared with the null model (*B*_10_ > 100), the model with “stage+site+stage×site” was the best and far outperformed the other models. The significant interaction between stage and site, and the best model with the interaction factor “stage×site”, revealed that the HRV and ear-PRV patterns across sleep stages were significantly different. These results implied that although the DFA1 for HRV and PRV were different, the pattern of finger-DFA1 across the sleep stages was much more similar to that of heart-DFA1 than that of ear-DFA1. However, although it was significantly different from heart-DFA1, the ear-DFA1 still provided weak but considerable information regarding the difference between the sleep stages (Table S4).

For the DFA2 of each recording site, the results showed the main effects of stage for HRV (*N* = 39, 12 women; mean age, 22.6 ± 3.6 years; *F*(4, 152) = 45.69;*p* < 0.00;, 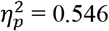, GG-corrected), finger-PRV (*N* = 25, 8 women; mean age, 22.0 ± 1.3 years; *F*(4, 96) = 19.51;*p* < 0.001; 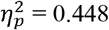, GG-corrected), and ear-PRV (*N* = 23, 6 women; mean age, 22.7 ± 4.6 years; *F*(4, 88) = 18.81; *p* < 0.001; 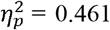, GG-corrected). The results of the comparisons of the patterns of the three recording sites across five different stages showed the main effects of stage (*N* = 12, 3 women; mean age, 21.7 ± 1.3 years; *F*(4, 44) = 20.66;*p* < 0.001; 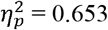) and site (*F*(2, 22) = 51.54;*p* < 0.001; 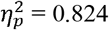, GG-corrected) However, the interaction between the two factors was not significant (*F*(8, 88) = 2.23; *p* = 0.090; 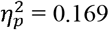, GG-corrected). The Bayes factors for model comparison (see Table S5) showed that compared with the null model (*B*_10_ > 100), the model with “stage+site” was the best and far outperformed the other models. The non-significant interaction between stage and site, and the best model without the interaction factor “stage×site”, revealed the same patterns for heart-DFA1, finger-DFA1, and ear-DFA1 across sleep stages.

Except for ApEn that showed a trend of difference, the non-linear finger and ear indices (DFA1 and DFA2) showed very similar patterns to those of the heart indices. Specifically, ApEn for HRV and finger-PRV were lower in NREM sleep (N1 and N2) than in Wk, and lower in N2 than in REM sleep. However, in NREM sleep, evidence showed that finger-ApEn was higher in N3 than in N2 and N1, which is opposite to the result for heart-ApEn. DFA1 and DFA2 showed similar results. For all three recording sites, DFA1 and DFA 2 were lower in NREM sleep (N2 and N3) than in REM sleep and Wk. In NREM sleep, they were lower in N3 than in N2 and N1, and in N2 than in N1. In contrast to DFA1, DFA 2 showed lower values in N1 than in Wk. However, the ear-PRV indices showed weaker evidence.

### 3.2 Comparison of the changes in 0.2–0.3 Hz oscillations in PRV and HRV power spectra across different sleep stages

As mentioned in the Methods section, we also investigated the (normalized) power spectra of the IBI signals derived from the BF and ECG signals (IBI_Pow_02_03). For the IBI_Pow_02_03 of HRV/PRV, which was normalized by dividing it by the HF(see Figure S1 and Table S4 for more information on the LFn, HFn, and LF/HF of HRV/PRV) for each recording site, the results (Figure 6) showed the main effects of stage for HRV (*N* = 40, 12 women; mean age, 22.6 ± 3.5 years; *χ*^2^(4) = 61.62; *p* < 0.001), finger-PRV (*N* = 28, 8 women; mean age, 22.1 ± 1.4 years; *F*(4, 108) = 3.95; *p* = 0.012; 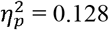, GG-corrected), and ear-PRV (*N* = 21, 6 women; mean age, 22.6 ± 4.7 years; *F*(4, 80) = 2.45; *p* = 0.081; 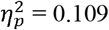 GG-corrected). Similarly, the data for IBI_Pow_02_03 for HRV in N3 violated normality; thus, it was subjected to the non-parametric two-way analysis using R with nparLD.[2,3] The results showed the main effects of stage (*N* = 14, 3 women; mean age, 21.6 ± 1.2 years; *χ*^2^(4) = 11.40; *p* = 0.022) and site *χ*^2^(2) = 41.94, *p* < 0.001). However, the interaction between the two factors was non-significant *χ*^2^(8) = 8.25, *p* = 0.410). The patterns for HRV, finger-PRV, and ear-PRV across sleep stages were all similar. For HRV, finger-PRV, and ear-PRV, normalized IBI_Pow_02_03 was higher in NREM (N2 and N3) sleep than in REM sleep; however, the evidence for ear-PRV was weak. For NREM sleep, IBI_Pow_02_03 was higher in N3 than in N1, and in N2 than in N1, for both HRV and finger-PRV; however, the evidence for finger-PRV was weaker.

**Figure 6.**
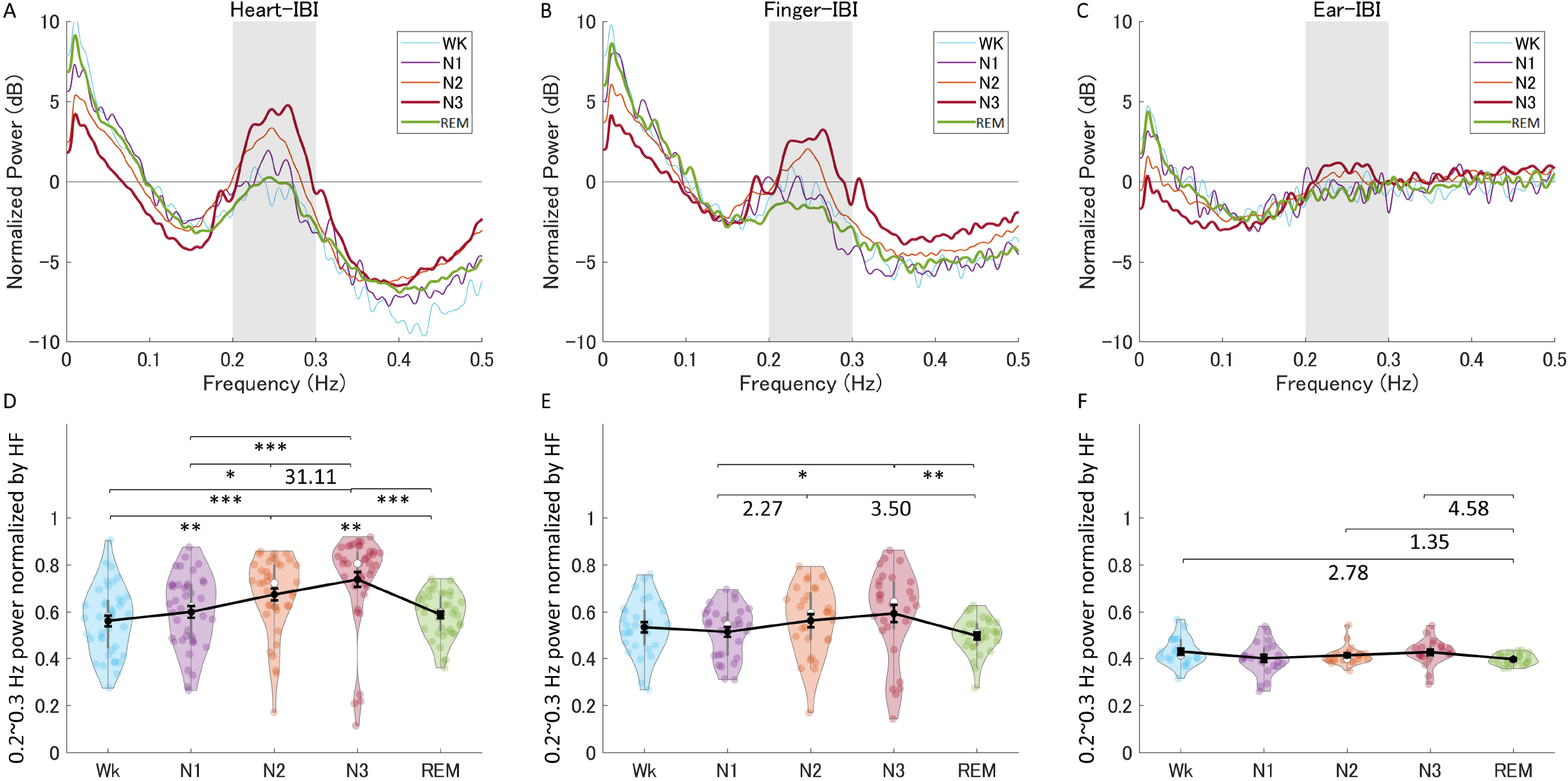
The normalized power spectra and the normalized power of the 0.2–0.3 Hz band of HRV, finger-PRV, and ear-PRV across the different sleep stages. (A-C) The normalized power spectra for HRV (A), finger-PRV (B), and ear-PRV (C). (D-F) The normalized power of the 0.2–0.3 Hz band for HRV (D), finger-PRV (E), and ear-PRV (F). There was a linear trend of power increase in the 0.2–0.3 Hz band with the deepening of sleep from N1 to N3 for finger-PRV, ear-PRV, and HRV. The violin plot with dots shows the distribution of the individual data points. The line chart with error bars shows the group mean and the ± 1 standard error of the mean. \**p* < 0.05; \*\**p* < 0.01; \*\*\**p* < 0.001. The numerical values represent the Bayes factors, which show anecdotal (1–3), moderate (3–10), or strong evidence (> 10) against the H0 of no difference between pairs of sleep stages. Bayes factors < 1 are not listed. HRV, heart rate variability; PRV, pulse rate variability; IBI, inter-beat interval; HF, high frequency power.

Regarding the same 0.2–0.3 Hz band power from IBI signals, finger-IBI showed consistency with heart-IBI, with N3 having the highest peak power. Trend analysis revealed a linear trend for finger-IBI (*p* < 0.001) and ear-BF (*p* = 0.010); however, ear-IBI provided weaker evidence.

### 3.3 Comparison of the changes in the time-domain parameters of finger- and ear-BF signals across different sleep stages

In addition to the PRV indices, we also investigated the time-domain and frequency-domain (in the following paragraphs) parameters of finger-BF and ear-BF, expecting that BF signals would provide additional information about the different sleep stages. Figure 7 shows the indices of finger-BF and ear-BF in the time domain, including the mean amplitude, SD, and CV of BF (Table S4). For the mean amplitude of each recording site (finger or ear), the results showed the main effects of stage for finger-BF (*N* = 28, 8 women; mean age, 22.1 ± 1.4 years; *F*(4, 108) = 3.61; *p* = 0.022; 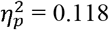, GG-corrected), but not for ear-BF (*N* = 22, 7 women; mean age, 23.0 ± 4.7 years; *F*(4, 84) = 2.13; *p* = 0.117; 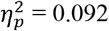, GG-corrected). For SD, the results showed the main effects of stage for finger-PRV (*N* = 27, 8 women; mean age, 22.1 ± 1.4 years; *F*(4, 104) = 3.93; *p* = 0.010; 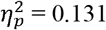, GG-corrected), but not for ear-PRV (*N* = 23, 7 women; mean age, 22.9 ± 4.6 years; *F*(4, 88) = 1.63; *p* = 0.193; 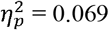, GG-corrected). For CV, the results showed the main effects of stage for finger-PRV (*N* = 25, 8 women; mean age, 22.0 ± 1.2 years; *χ*^2^(4) = 16.90; *p* = 0.002), but not for ear-PRV (*N* = 21, 7 women; mean age, 23.1 ± 4.8 years; *χ*^2^(4) = 3.28; *p* = 0.513).

**Figure 7.**
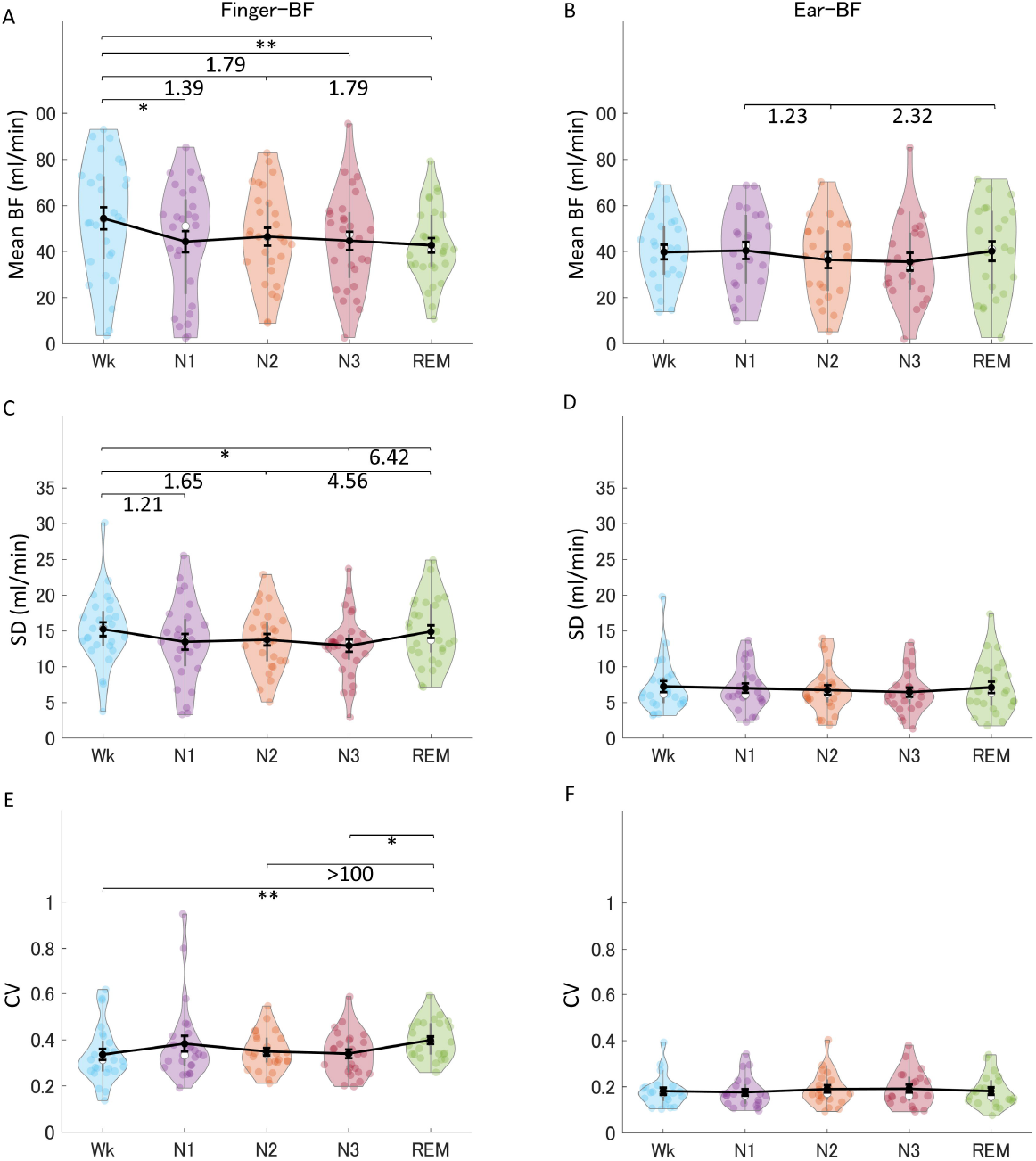
The indices of finger-BF and ear-BF in the time domain, including the mean (A-B), SD (C-D), and CV (E-F) of BF, across the different sleep stages. The violin plot with dots shows the distribution of the individual data points. The line chart with error bars shows the group mean and the ± 1 standard error of the mean. \**p* < 0.05; \*\**p* < 0.01. The numerical values are the Bayes factors. Only those for non-significant comparisons were provided here for complementary information (for other information, see Table S4). The values show anecdotal (1–3), moderate (3–10), strong (> 10), or extreme evidence (> 100) against the H0 of no difference between pairs of sleep stages. Bayes factors < 1 are not listed. BF, blood flow; SD, standard deviation; CV, coefficient of variance.

The results revealed that finger-BF was modulated differently across sleep stages. Interestingly, finger-BF showed a robust difference in mean amplitude and CV between Wk and REM sleep. All other indices of finger-BF and PRV did not show a difference between the two stages, suggesting that BF signal itself may provide irreplaceable information about the sleep stages compared with PRV. Interestingly, the mean finger-BF in Wk was larger than that in the REM stage, whereas CV, reflecting the relative fluctuation, was larger in REM than in Wk. However, except for some weak evidence, ear-BF was not significantly modulated by the sleep stages.

### 3.4 Comparison of the changes in 0.2–0.3 Hz oscillations of BF spectra across different sleep stages

Figures 8A and B show the power spectra of finger- and ear-BF. Interestingly, we noted a peak power within the HF band (0.15–0.4 Hz), specifically around 0.2–0.3 Hz. Furthermore, the 0.2–0.3 Hz band power was modulated by the sleep stages. Figures 8D and E show the power in the 0.2–0.3 Hz band (referred to as BF_Pow_02_03; see the dark area in the figure) across sleep stages. In addition, a representative BF signal modulated in a 0.2–0.3 Hz oscillation during N3 is shown in Figure 9.

**Figure 8.**
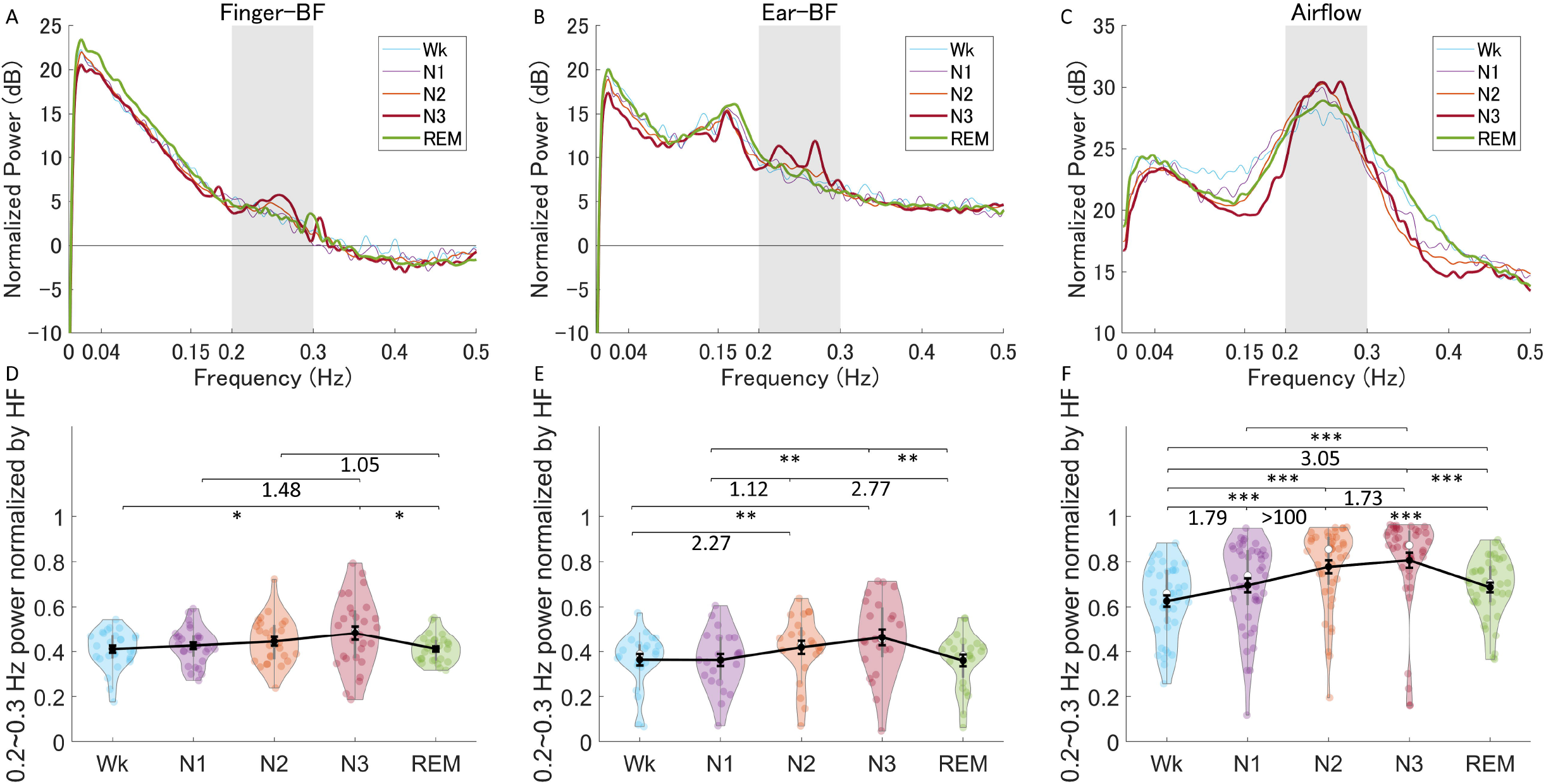
The normalized power spectra and the normalized power in the 0.2–0.3 Hz band of finger-BF, ear-BF, and AF, across the different sleep stages. (A-C) The normalized power spectra for finger-BF (A), ear-BF (B), and AF (C). (D-F) The normalized power of the 0.2–0.3 Hz band for finger-BF (D), ear-BF (E), and AF (F). There was a linear trend of power increase in the 0.2–0.3 Hz band with the deepening of sleep from N1 to N3 for both finger- and ear-BF. The violin plot with dots shows the distribution of the individual data points. The line chart with error bars shows the group mean and the ± 1 standard error of the mean. \**p* < 0.05; \*\**p* < 0.01. The numerical values are the Bayes factors. Only those for non-significant comparisons were provided here for complementary information (for other information, see Table S4). The values show anecdotal (1–3) evidence against the H0 of no difference between pairs of sleep stages. Bayes factors < 1 are not shown. BF, blood flow; HF, high frequency power.

**Figure 9.**
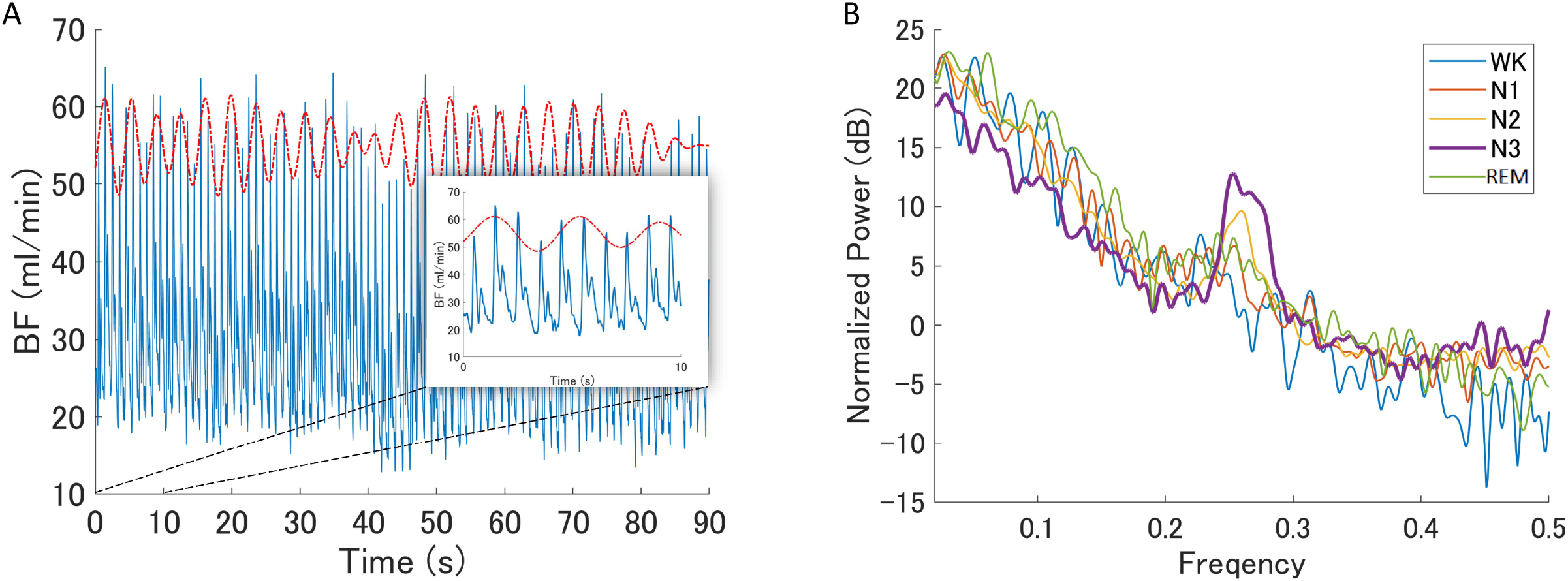
BF modulated by oscillations within the 0.2–0.3 Hz band. (A) A representative epoch of the raw BF signal modulated by 0.2–0.3 Hz oscillations during N3 recorded from a representative participant. The red dotted curve represents the oscillating signal filtered from the raw BF signal with a bandpass frequency of 0.2–0.3 Hz; however, it has been shifted upwards to make it easier to read. The blue curve is the raw BF signal. (B) The normalized power spectrum of a representative participant at different sleep stages. It shows a peak in the 0.2–0.3 Hz frequency band during N3. BF, blood flow.

For the 0.2–0.3 Hz band power of BF, which was normalized by dividing it by the HF(see Figure S1 and Table S4 for more information on LFn, HFn, and LF/HF of BF) for each recording site, the results showed the main effects of stage for finger-BF (*N* = 28, 8 women; mean age, 22.1 ± 1.4 years; *F*(4, 108) = 3.57; *p* = 0.018; 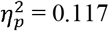, GG-corrected) and ear-BF (*N* = 23, 6 women; mean age, 22.7 ± 4.6 years; *F*(4, 88) = 4.98; *p* = 0.004; 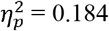, GG-corrected). For both finger- and ear-BF, the normalized BF_Pow_02_03 was higher in NREM (N3) sleep than in REM sleep and Wk. In NREM sleep, BF_Pow_02_03 was higher in N3 than in N1; however, the evidence for finger-BF was weak. We then extracted the data for N1–N3 and conducted a trend analysis, which is one of the sub-analyses in the F-RMANOVA in JASP. The results showed a linear trend in finger-BF (*p* = 0.001) and ear-BF (*p* = 0.001). Finger-BF was consistent with ear-BF with respect to BF_Pow_02_03 across the NREM sleep stages N1– N3; both had the highest value in N3.

### 3.5 Comparison of the changes in 0.2–0.3 Hz oscillations of AF spectra across different sleep stages and their correlation with BF

We have observed special 0.2–0.3 Hz oscillations in BF, which is within the respiratory frequency band of 0.15–0.4 Hz. This band of oscillations may be modulated by respiration activity. Therefore, we analyzed the AF data and investigated the correlation between the BF and AF activities (see the supplementary information for the methods) to provide more insight into the mechanism underlying the 0.2–0.3 Hz oscillations in BF. Figures 8C and F show the power spectra of AF and the power in the 0.2– 0.3 Hz band (referred to as AF_Pow_02_03; see the dark area in the figure), which was normalized by dividing it by the HF(see Figure S1 and Table S4 for more information on the LFn, HFn, and LF/HF of AF). The results showed the main effect of stage (*N* = 41, 12 women; mean age, 22.5 ± 3.5 years; *χ*^2^(4) = 56.82; *p* < 0.001). AF_Pow_02_03 also seemed to be a good indicator for distinguishing between sleep stages. Further, correlations between the 0.2–0.3 Hz power and the peak frequency were significant in several sleep stages, especially in N2 and N3, suggesting a close relationship between peripheral BF and respiration (see Figure S2).

## 4 Discussion

In this study, we investigated BF and PRV across different sleep stages and compared PRV indices with those of HRV during sleep. Comparisons of the time-domain, frequency-domain, and non-linear PRV/HRV indices across the sleep stages revealed that the patterns of finger-PRV indices were consistent with those of most HRV indices, especially for the time-frequency domain. However, differences existed between finger-PRV and HRV indices, especially in non-linear metrics such as ApEn. Ear-PRV could also provide information for differentiating sleep stages, comparable to the information provided by some of the HRV indices. In addition, the time- and frequency-domain parameters of BF signal could also provide some information regarding the different sleep stages.

We observed significant peaking of oscillations around 0.2–0.3 Hz, especially during N3, for both BF signal and the derived IBI signal. For finger- and ear-BF, finger- and ear-PRV, and HRV, we noted an increase in oscillations within the 0.2–0.3 Hz band with the deepening of NREM sleep (N1 to N3). This increase was highest in N3. The findings of this study suggest that PRV can provide as much information as HRV about the sleep state. In some respects, BF+PRV may provide more information than HRV alone, e.g., 0.2–0.3 Hz oscillations during different sleep stages. Therefore, the potential of BF+PRV to provide information in the sleep state requires further investigation.

### 4.1 Peripheral BF dynamics across sleep stages

Normal human sleep is associated with hemodynamic changes, which are primarily mediated by changes in the ANS. Specifically, during NREM sleep, sympathetic activity decreases, and parasympathetic (vagal) activity increases with the development of slow-wave sleep (N3). In contrast, there is a reversal of these two changes during REM sleep.[43,44] These distinct changes during different sleep stages are reflected in heart rate and HRV/PRV. In the present study, the mean IBI, an index of changes in the balance of the two ANS components, increased (i.e., heart rate decreased) during NREM sleep with the sleep deepening that occurs from N1 to N3. However, it decreased during REM sleep (more specifically, phasic REM sleep) but remained above Wk.[43,45,46] The mean finger- and ear-IBIs in the present study showed differences between nearly all pairs of different sleep stages except for N1 and REM sleep.

PRV indices are also modulated by sleep stages with different ANS activity patterns. Frequency-domain indices, including LFn, HFn, and LF/HF, showed nearly the same patterns across sleep stages as the mean IBI. They also showed differences between most pairs of different sleep stages, except between Wk, N1, and REM sleep. Although LF reflects a mix of sympathetic and parasympathetic power and HF reflects vagal tone, the calculation of LFn, HFn, and LF/HF makes them equivalent in bearing information on the patterns of ANS activity across sleep stages.[3,16] Time-domain indices, including pNN50, are correlated with RMSSD, as well as with HF; therefore, the two indices in the time domain (for finger-PRV) showed similar patterns. In contrast, LF makes a significant contribution to SDNN; thus, the SDNN pattern for finger-PRV was similar to that of LFn.[3] The non-linear indices of both finger- and ear-PRV, specifically DFA1 and DAF2, measure the fluctuations of IBIs. Thus, it is not surprising that they bear as much information for differentiating sleep stages as the frequency-domain indices that measure the oscillations of IBIs. The patterns of HRV/finger-PRV indices (including SDNN, LFn, HFn, and LF/HF) were consistent with a recent study.[12] However, that study did not provide significant test results for the comparisons between the different sleep stages.

Besides PRV, BF may be modulated differently by the ANS during different sleep stages. For example, the peripheral arterial tone of BF, which is an index of sympathetic vasoconstrictor mechanisms,[47] decreases from Wk to NREM sleep, and reaches nadir during REM sleep.[48] The decreasing pattern of the peripheral arterial tone suggests a similar decreasing pattern for BF across sleep stages because they share similarities in the assessment of peripheral pulse waves.[49] The mean amplitude of BF in the present study confirmed this; however, the evidence was weak. In addition, because REM sleep is associated with a largely variable sympathetic tone,[47] BF during this stage should be variable as well. This was also confirmed by the CV results of BF in the present study. Furthermore, previous studies have shown that the modulation of BF by sleep stages (e.g., REM sleep with higher sympathetic activity) may be different between the peripheral and cerebral regions. While peripheral BF may decrease in REM sleep compared with NREM sleep,[48] cerebral BF shows a marked increase instead.[46,50] Therefore, besides PRV, which can be compared with HRV, peripheral BF has its own characteristics during sleep, and these characteristics may be different across sleep stages. BF+PRV not only differentiate the sleep stages, but also make good indices for the measurement of ANS activity, including parasympathetic and sympathetic. In some respects, they compare favorably with HRV because HRV indices (e.g., LFn) do not have mono-measures for sympathetic tone. However, BF indices, such as mean and CV, may be candidates for the assessment of sympathetic nervous system activation.

### 4.2 The similarities and differences between PRV and HRV regarding the ANS activity during sleep

Anatomically, BF and ECG signals are closely related. With every heartbeat, blood is transferred from the heart through the blood vessel network to peripheral areas, such as fingers. Because of this close relationship, attempts have been made in previous studies to reconstruct ECG signals from the pulse wave signals of BF.[51] Although many external factors can affect the consistency between PRV and HRV, the physiological nature of BF and ECG signals leads to inevitable differences between them. One critical intrinsic factor is the pulse transit time, which is the time required for the blood to travel from the heart to the peripheral site where the BF is measured. Therefore, it is expected that traveling through different blood vessel networks to different sites, such as the finger and ear, may affect the PPT and could affect the signals recorded and the information they provide. In addition, the two components of the ANS, the parasympathetic and sympathetic, have different roles in the heart and vasculature.[17] While the sympathetic nervous system plays a dominant role in regulating vascular activity, the parasympathetic nervous system majorly contributes to cardiac activity. For example, many HRV indices, such as RMSSD, pNN50, and HF, reflect vagal tone. Other HRV indices, such as LF, reflect a mix of vagal and sympathetic activity,[16] suggesting a major role of vagal tone in HRV. However, BF and PRV may be affected by factors that affect HRV through the blood vessel network, and by factors that directly affect vascular activity, such as the sympathetic system[17,18] or changes in the local vasculature.[8.9] Thus, variations in BF and PRV can come from different sources that do not affect HRV in the same way. This may also explain why some indices, such as ApEn, showed a trend of difference between PRV and HRV. This may be because PRV may be more complex than HRV, even during N3. The differences between PRV and HRV require further detailed investigation.

### 4.3 The 0.2–0.3 Hz oscillations during sleep

HFn (0.15–0.40 Hz) reflects respiratory sinus arrhythmia used to measure parasympathetic (vagal) activity. The 0.2–0.3 Hz oscillations are within the HFn frequency band. This frequency band is consistent with the so-called Traube-Hering waves, which refer to blood pressure oscillations in time with breathing.[52] A plausible explanation for this is that the band may reflect the respiratory modulation of BF and HRV/PRV. Both finger- and ear-BF showed an apparent peak power in 0.2–0.3 Hz, which increased with the deepening of sleep, e.g., from N1 to N3. In addition, the relative power in the 0.2–0.3 Hz band of both finger- and ear-BF correlated with that of respiratory activity. Previous studies have also reported that respiratory events modulate the pulse wave amplitude of BF more than ECG/HRV.[11,53] PRV can reflect the coupling effect between respiration and vasculature. However, although there have been studies on the effects of respiration around 0.2–0.3 Hz on cardiovascular activity,[7,35] no studies have been conducted to investigate this band of oscillations during sleep, especially by comparing it among sleep stages. The results of the present study showed that oscillations occur in the 0.2–03 Hz band during sleep and that sleep stages modulate these oscillations. Interestingly, the 0.2–0.3 Hz oscillations were largest during N3 (deep sleep) and were reflected in BF, PRV and HRV.

Sleep exerts effects on breathing patterns and ANS activity, and thus, on BF and PRV. For example, during N3, both BF and PRV showed the 0.2–0.3 Hz oscillations, and the 0.2–0.3 Hz oscillations of BF were correlated with those of AF. Conversely, changes in breathing patterns and ANS activity may modulate sleep. Supposing the 0.2–0.3 Hz oscillations can be affected by external stimulation, such as sensory stimulation (e.g., rocking), sleep may be modulated by this pathway. A recent study conducted using 0.25 Hz rocking stimulation successfully entrained spontaneous neural oscillations with benefits for sleep and memory.[32] The present study provides consistent evidence for using stimulation of approximately 0.25 Hz to facilitate sleep. However, to the best of our knowledge, the causal effects of stimulation in the 0.2–0.3 Hz band on BF activity during sleep has not been investigated yet; thus, future studies needed to clarify this.

### 4.4 Determining different sleep stages using BF+PRV

Compared with ECG/HRV, BF/PRV can be measured in a much more convenient, straightforward, low-cost, and non-invasive way, and can be measured at more recording sites. PRV has been used to predict sleep stages in previous studies with acceptable accuracy [54]. However, the BF signal itself is often ignored, and only some of the PRV features are used, which may be the reason for the lack of high accuracy. Some researchers have also tried to utilize both the BF or the corresponding photoplethysmography signal and PRV to automatically score sleep stages. However, algorithms were used in these studies to extract features that cannot be easily related to the physiological origins of the sleep stages.[55] The present study may have provided new features for determining different sleep stages and understanding their physiological origins.

### 4.5 Limitations of this study

Although measurement of PRV is much more convenient, factors that affect PRV rather than HRV include noise in the signal, errors in detecting the fiducial points, and physiological factors such as the pulse transit time and changes (or respiration factors leading to changes) in blood pressure. The noise mainly comes from unstable attachment of the sensors to body areas (e.g., ear concha) and is related to body movement. With further development of these sensors, the noise problem may be solved. Thus, noise during data acquisition was a drawback in the present study. Several measurement data and a certain number of epochs from individual data were excluded from analysis because of noise during acquisition (Table S1) and interruptions during recording. The right-index finger and right-ear concha sites are easily affected by body movements and posture. Thus, the first limitation of this study was the movement artifact that affects the quality of BF data. Future device designs should focus on increasing the signal-to-noise ratio. Owing to the noise, the quality of fiducial point detection was affected. The second limitation of this study was the evaluation of the systolic peaks for the detection of the fiducial point for PRV as an analogy to the actions of R peaks for HRV. However, this method is reported to show the lowest agreement between PRV and HRV. This may not greatly affect this study because we found that finger-PRV showed much consistency with HRV. However, ear-PRV must be significantly affected. Even during preprocessing, we discovered that detection of the fiducial point was complex for many epochs of ear-BF because of noise. Therefore, it may not be intrinsic factors, such as the nature of BF and ECG, which affect the consistency between the two, but may instead be external factors such as technical design and noise, including movement artifacts that affect the application of BF+PRV. The third limitation is that we only recruited healthy participants in this study. No patients, e.g., those with sleep apnea, were included. Thus, this study cannot be directly compared with previous studies of patients.[11,12] The fourth limitation is that we only conducted the experiment during nighttime sleep rather than during a daytime nap. HRV is modulated by the circadian rhythm,[56] which suggests that the dynamics of BF and PRV during sleep stages shown in this study may differ between a daytime nap and nighttime sleep. Nevertheless, the present study has proved that BF+PRV provides much information about sleep stages, which may be more than that provided by HRV alone, supporting the prospect of BF+PRV as a new biomarker.[15] However, this conclusion should be explored in future studies.

In conclusion, the BF signal and the derived IBI signal provide considerable information about the sleep state, specifically, the sleep stages. Time-domain and frequency-domain BF parameters, and time-domain, frequency-domain and non-linear PRV indices provide ample information about the sleep stages, possibly more than HRV alone. BF+PRV should be considered a new biomarker instead of a surrogate of ECG/HRV. Its potential for monitoring and predicting sleep state should be fully explored.

## 5 Acknowledgments

This study was supported by the KYOCERA Corporation of Japan.

## 7 Data Availability Statement

The data underlying this article cannot be shared publicly due to a non-disclosure agreement with KYOCERA Corporation of Japan. The data underlying this article will be shared on reasonable request to the corresponding author.

