## Supplementary information for "Dynamics of peripheral blood flow across sleep stages"

Institution where work was performed: International Institute for Integrative Sleep Medicine (WPI-IIIS), University of Tsukuba, Tsukuba, Japan.

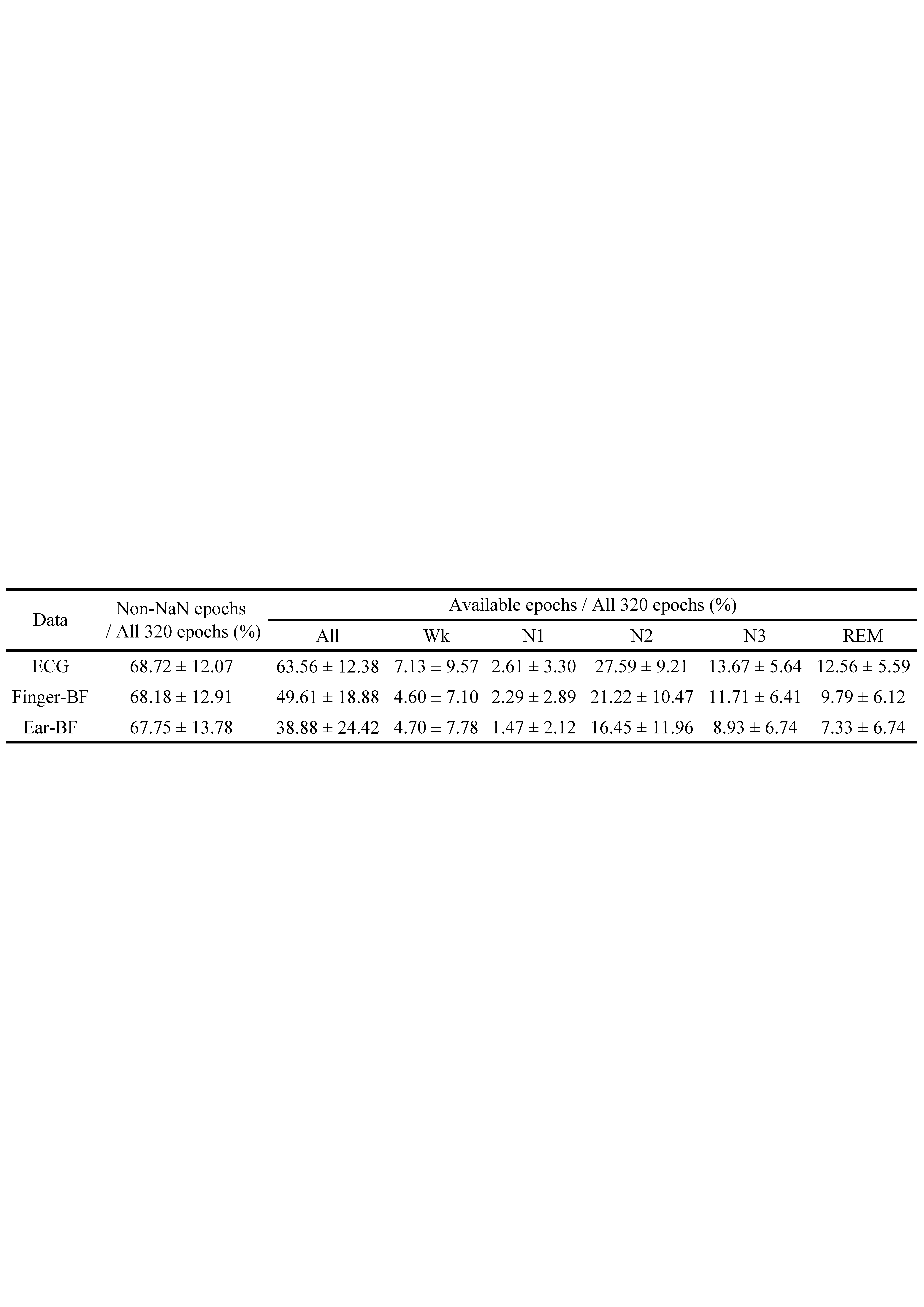
**Table S1. Usability of data (epochs) for ECG, finger-BF, and ear-BF across the sleep stages**

Note: The percentage here is the average of all available individual percentages.

NaN, not a number; Wk, wakefulness; N1–3, non-rapid-eye-movement sleep 1–3; REM, rapid-eye-movement sleep; ECG, electrocardiogram; BF, blood flow.

**Table S2. PRV/HRV indices selected for analysis**

| MeanIBI | Mean of inter-beat intervals corresponding to R-to-R intervals or pulse-to-pulse of blood flow intervals |
| --- | --- |
| Time-domain indices | |
| SDNN | Standard deviation of all the normal IBIs (normal-to-normal [NN] intervals) |
| RMSSD | Root mean square of successive differences between the adjacent NN intervals |
| pNN50 | Percentage of pairs of the adjacent NN intervals differing by more than 50 ms |
| Frequency domain indices | |
| LFn | Normalized low frequency (0.04–0.15 Hz) power: LF/(LF+HF) |
| HFn | Normalized high frequency (0.15–0.40 Hz) power: HF/(LF+HF) |
| LF/HF | Ratio of LF to HF |
| Non-linear measurements | |
| ApEn | Approximate entropy |
| DFA1 | Detrended fluctuation analysis, which measures short-range fluctuations (4 to 12 beats) |
| DFA2 | Detrended fluctuation analysis, which measures long-range fluctuations (13 to 64 beats) |

Note: PRV, pulse rate variability; HRV, heart rate variability; R, R peaks; IBI, inter-beat intervals; LF, low frequency power; HF, high frequency power.

**Table S3. Evidence categories for the Bayes factor** $\boldsymbol{B}_{\boldsymbol{10}}$

| Bayes factor $B_{10}$ | Interpretation |
| --- | --- |
| > 100 | Extreme evidence for H1 |
| 30 – 100 | Very strong evidence for H1 |
| 10 – 30 | Strong evidence for H1 |
| 3 – 10 | Moderate evidence for H1 |
| 1 – 3 | Anecdotal evidence for H1 |
| 1 | Insufficient evidence for either H1 or H0 |
| 0.33 – 1 | Anecdotal evidence for H0 |
| 0.1 – 0.33 | Moderate evidence for H0 |
| 0.03 – 0.1 | Strong evidence for H0 |
| 0.01 – 0.03 | Very strong evidence for H0 |

Note: H1, the alternative hypothesis; H0, the null hypothesis; $B_{10}$, Bayes factor for H1 to H0.

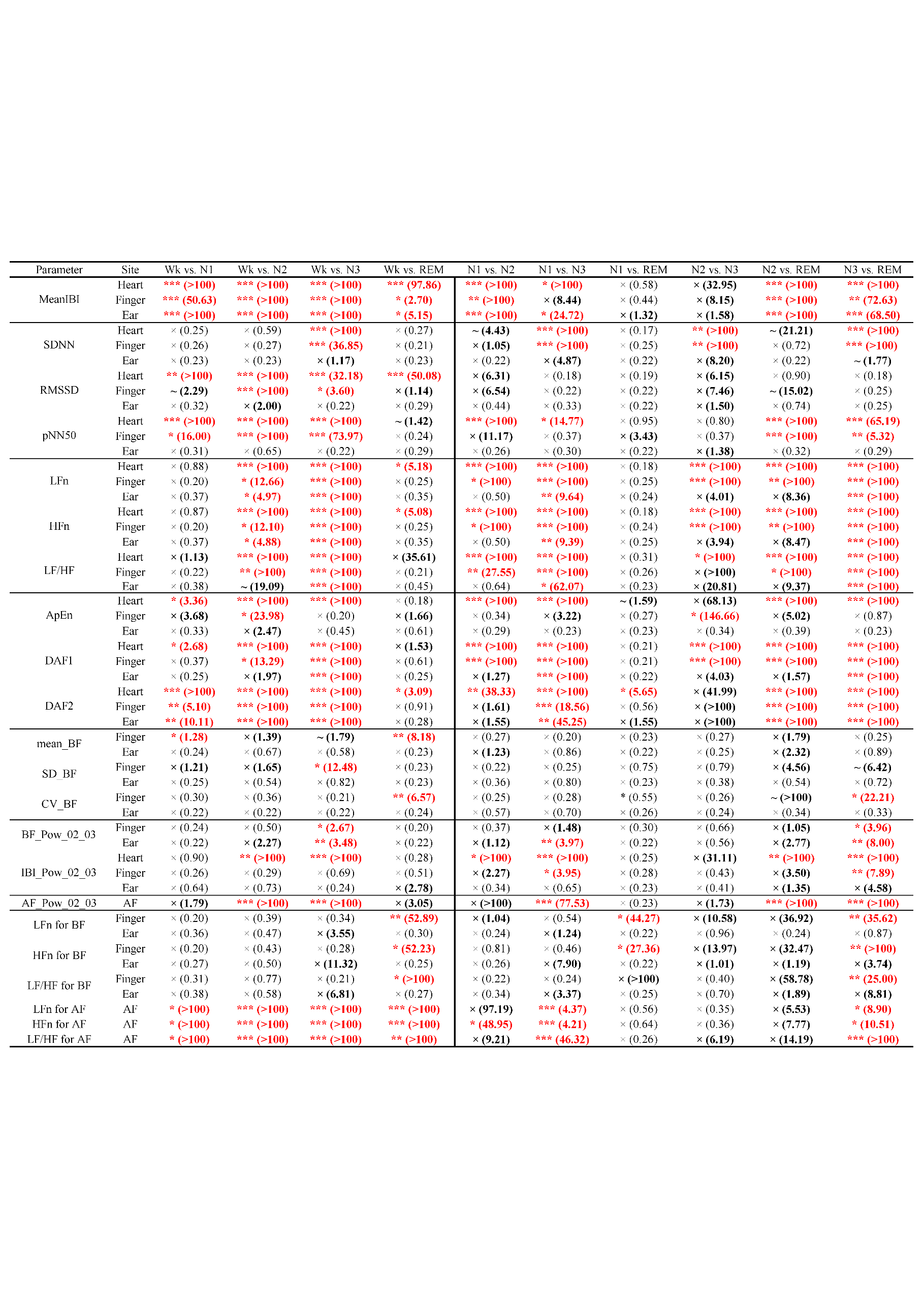
**Table S4. Significance test results and Bayes factors of different pairs of sleep stages for the HRV/PRV/BF indices measured at each recording site**

~: 0.05 < *p* <0.1; **p* < 0.05; ***p* < 0.01; ****p* < 0.001; ×non-significant

The numerical values are the Bayes factors. Values > 1 show that the evidence supports H1, values < 1 indicate that the evidence supports H0, and values equal to 1 indicate insufficient evidence supporting H1 or H0.

Note: HRV, heart rate variability; PRV, pulse rate variability; BF, blood flow; Wk, wakefulness; N1–3, non-rapid-eye-movement sleep 1–3; REM, rapid-eye-movement sleep; MeanIBI, mean of inter-beat intervals corresponding to R-to-R intervals or pulse-to-pulse of blood flow intervals; SDNN, standard deviation of all the normal-to-normal intervals; RMSSD, root mean square of successive differences between the adjacent normal-to-normal intervals; pNN50, percentage of pairs of the adjacent normal-to-normal intervals differing by more than 50 ms; LFn, normalized low frequency power; HFn, normalized high frequency power; ApEn, approximate entropy; DFA, detrended fluctuation analysis; SD, standard deviation; CV, coefficient of variance; BF_Pow_02­_03, power of BF data in 0.2–0.3 Hz; IBI_Pow_02­_03, power of inter-beat intervals data in 0.2–0.3 Hz; AF, airflow; AF_Pow_02­_03, power of airflow data in 0.2–0.3 Hz.

**Table S5. The results of the significance test performed using F-RMANOVA and the model comparisons performed using B-RMANOVA for each PRV/HRV index across three recording sites and five sleep stages**

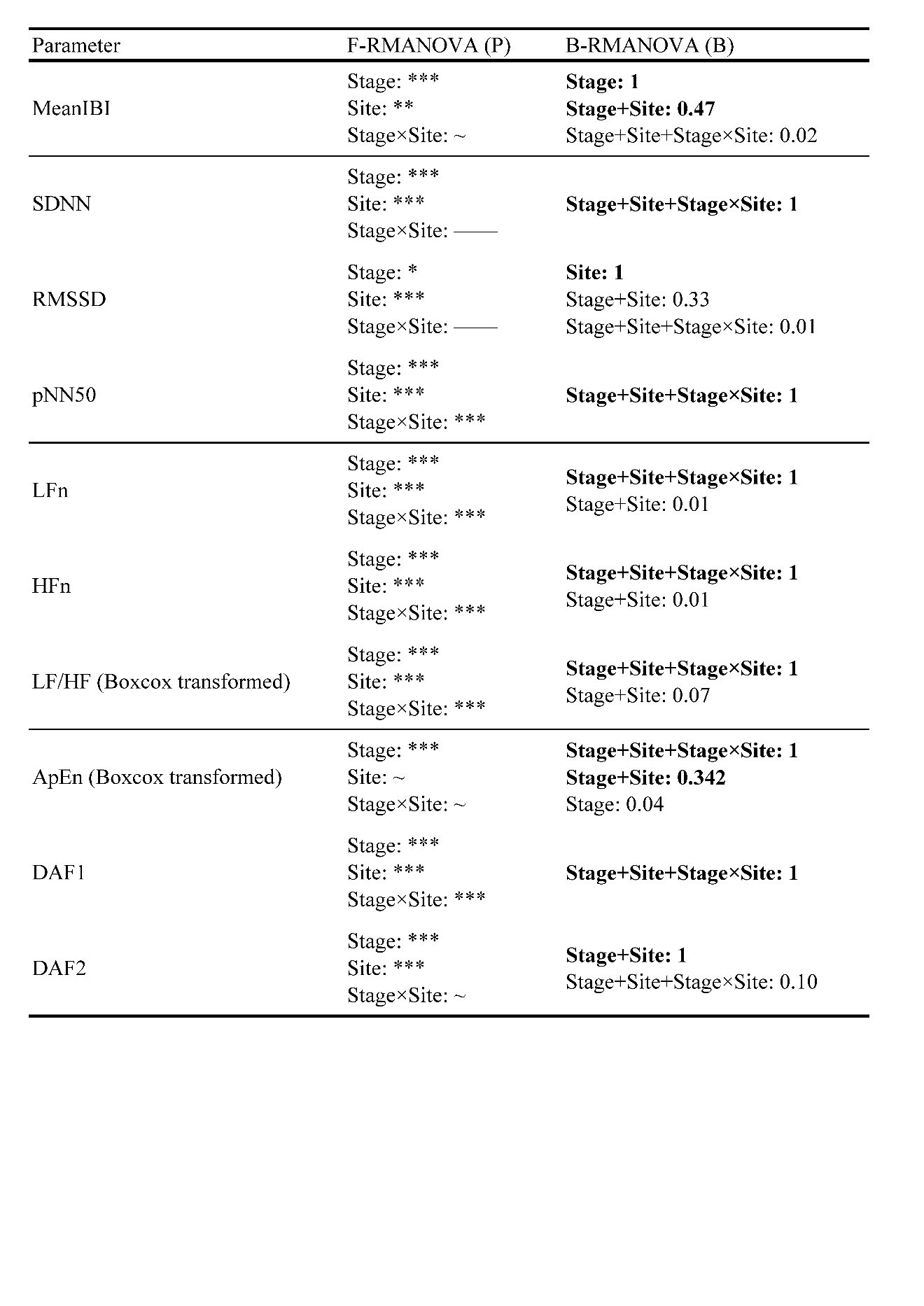

~0.05 < *p* <0.1; **p* < 0.05; ***p* < 0.01; ****p* < 0.001; ——non-significant

The numerical values are the Bayes factors, all of which were compared to the best model. Therefore, the best model always has a Bayes factor equal to 1. Models with Bayes factors < 0.01 (including the null model) are not shown. Thus, the best model is at least 100 times better than the null model.

Note: F-RMANOVA, frequentist repeated measure analysis of variance; B-RMANOVA, Bayesian repeated measure analysis of variance; PRV, pulse rate variability; HRV, heart rate variability; P, p value; B, Bayes factor; MeanIBI, mean of inter-beat intervals corresponding to R-to-R intervals or pulse-to-pulse of blood flow intervals; SDNN, standard deviation of all the normal-to-normal intervals; RMSSD, root mean square of successive differences between the adjacent normal-to-normal intervals; pNN50, percentage of pairs of the adjacent normal-to-normal intervals differing by more than 50 ms; LFn, normalized low frequency power; HFn, normalized high frequency power; ApEn, approximate entropy; DFA, detrended fluctuation analysis.

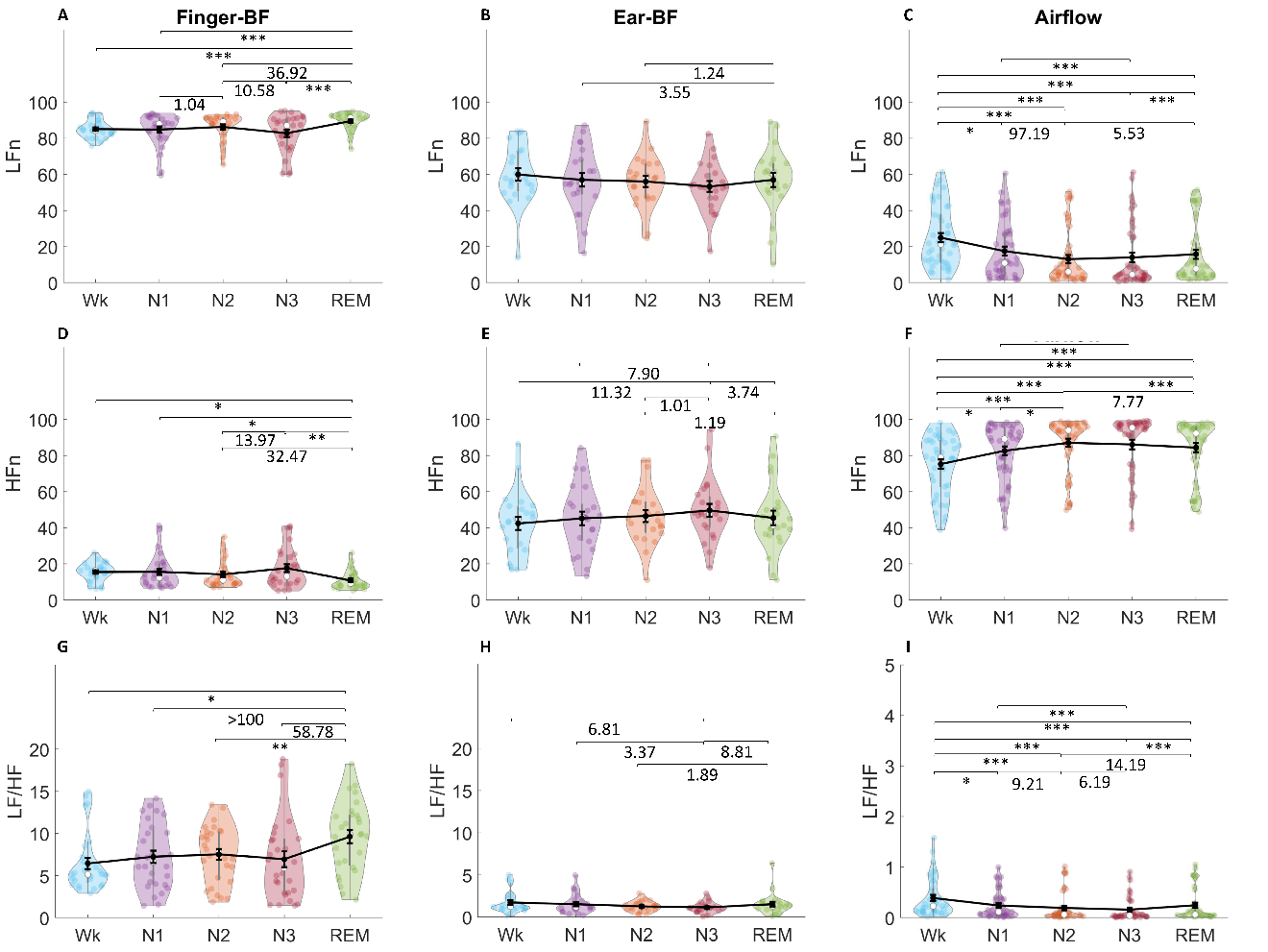

**Figure S1**. LFn (A, B, C), HFn (D, E, F), and LF/HF (G, H, I) of finger-BF (A, D, G), ear-BF (B, E, H), and AF (C, F, I) across the different sleep stages. The violin plot with dots shows the distribution of the individual data points. The line chart with error bars shows the group mean and the ± 1 standard error of the mean. The numerical values are the Bayes factors. Only those for non-significant comparisons were provided here for complementary information (for other information, see Table S4). The values show anecdotal (1–3), moderate (3–10), or strong (> 10) evidence against the H0 of no difference between the pairs of sleep stages. Bayes factors < 1 are not listed.

**p* < 0.05; ***p* < 0.01; ****p* < 0.001; LFn, normalized low frequency power; HFn, normalized high frequency power; LF, low frequency power; HF, high frequency power; BF, blood flow; AF, airflow.

Figure S1 shows the normalized low frequency power (LFn), normalized high frequency power (HFn), and low frequency power (LF)/high frequency (HF) power of finger-blood flow (BF), ear-BF, and airflow (AF) (see Table S4 for the significance test of the differences in LFn, HFn, and LF/HF between sleep stages for each site). For LFn, the results showed the main effects of stage for finger-BF (*N* = 27, 8 women; mean age, 22.1 ± 1.4 years; $\chi^{2}$(4) = 19.50; *p* < 0.001) and AF (*N* = 42, 12 women; mean age, 22.6 ± 3.5 years; $\chi^{2}$(4) = 65.56; *p* < 0.001), but not for ear-BF (*N* = 22, 6 women; mean age, 22.9 ± 4.7 years; *F*(4, 84) = 1.79; *p* = 0.165; $\eta_{p}^{2}$ = 0.079, GG-corrected).

For HFn, the results showed the main effects of stage for finger-BF (*N* = 27, 8 women; mean age, 22.1 ± 1.4 years; $\chi^{2}$(4) = 18.61; *p* < 0.001) and AF (*N* = 42, 12 women; mean age, 22.6 ± 3.5 years; $\chi^{2}$(4) = 65.81; *p* < 0.001), but marginally for ear-BF (*N* = 23, 7 women; mean age, 22.9 ± 4.6 years; $\chi^{2}$(4) = 9.15; *p* = 0.058).

For LF/HF, the results showed the main effects of stage for finger-BF (*N* = 28, 8 women; mean age, 22.1 ± 1.4 years; $\chi^{2}$(4) = 17.43; *p* = 0.002) and AF (*N* = 39, 11 women; mean age, 22.7 ± 3.6 years; $\chi^{2}$(4) = 68.70; *p* < 0.001), but not for ear-BF (*N* = 22, 7 women; mean age, 22.0 ± 1.4 years; $\chi^{2}$(4) = 8.07; *p* = 0.089).

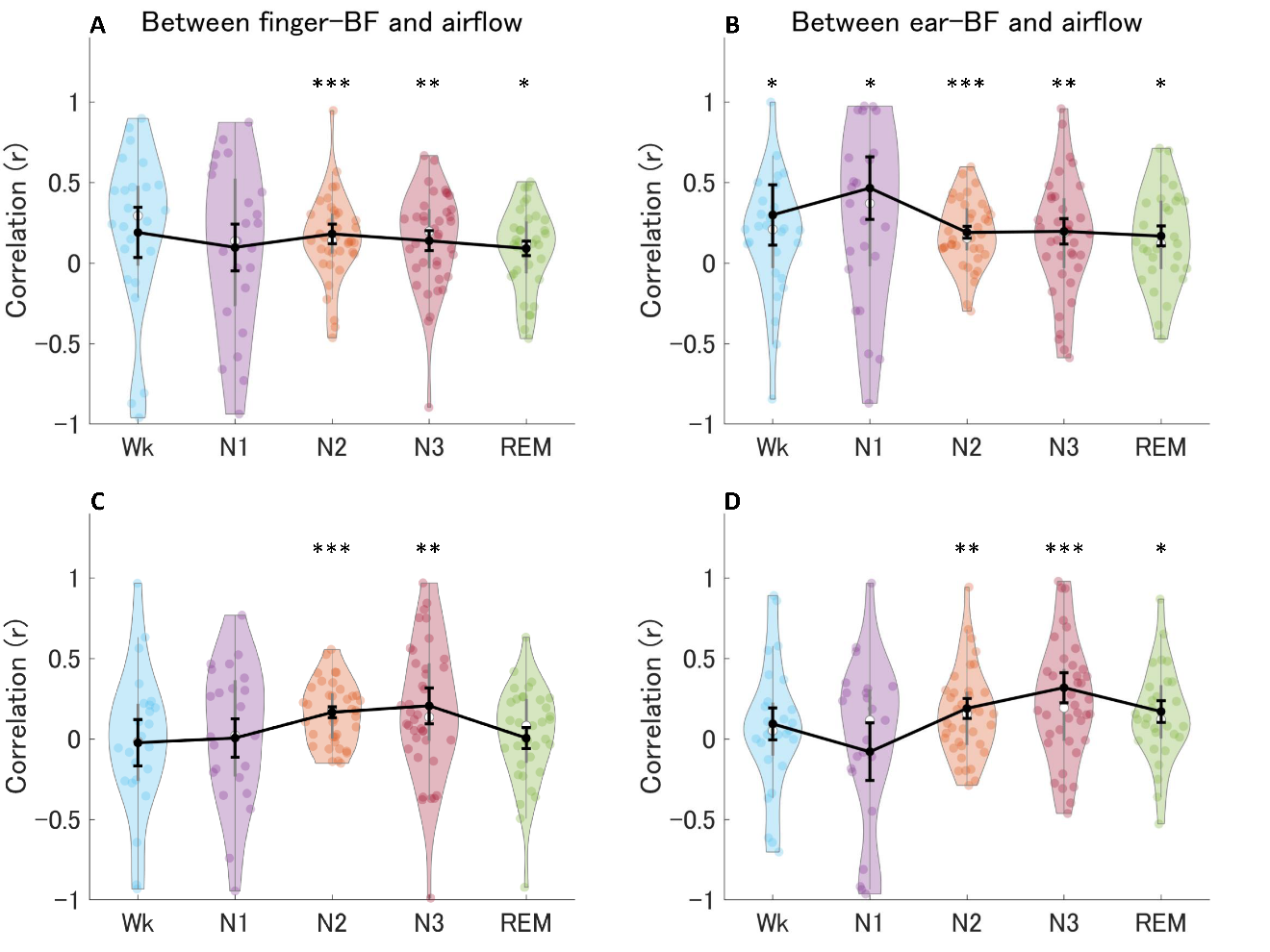

**Figure S2.** The correlations between the relative band power of 0.2–0.3 Hz for finger-BF and AF (A), and ear-BF and AF (B), as well as the correlations between the peak frequency within 0.2–0.3 Hz for finger-BF and the peak frequency within 0.15–0.4 Hz for AF (C), and ear-BF and AF (D). The violin plot with dots shows the distribution of the individual data points. The black line chart with error bars shows the inverse Fisher z-transformation of the group mean of Z coefficients (transformed from r) and the inverse Fisher z-transformation of the ± 1 standard error of the mean.

**p* < 0.05; ***p* < 0.01; ****p* < 0.001; BF, blood flow; AF, airflow.

**Methods used to obtain the results in Figures 8C and F, and in Figure S2**

Figure S2 shows the correlations between the relative band power of 0.2–0.3 Hz for finger-/ear-blood flow (BF) and airflow (AF), as well as the correlations between the peak frequency within 0.2–0.3 Hz for BF and the peak frequency within 0.150.4 Hz for AF, across epochs in each sleep stage.

Similar to the analysis of BF, 90-s epochs of AF from each individual were preprocessed using the customized program in MATLAB (The Mathworks Inc., Natick, MA, USA). The epochs with the largest amplitude beyond the three standard deviations of the median value of the highest amplitudes were excluded to remove epochs with significant artifacts. In addition, epochs with peaks of the waveform of AF (filtered using the default band-pass filter of 0–0.5 Hz embedded in the FieldTrip toolbox[22] beyond the three standard deviations of the median value of all the peaks were also excluded to ensure the quality of the beat signals was optimal.

The power spectra of AF was also analyzed using the “plomb” function in MATLAB. The band power of 0.2–0.3 Hz was extracted, normalized by dividing it by the high frequency power (as well as the normalized high- and low- frequency power, and low-/high- frequency power of AF, see Figure S1), and compared. The individual AF values larger than three standard deviations of the group means were set as missing values. Participants with missing values in any of the five stages were excluded from the group analysis.

The correlations between the relative band power of 0.2–0.3 Hz for BF and AF, and between the peak frequency within 0.2–0.3 Hz for BF and that within 0.15–0.4 Hz for AF, were calculated across epochs in each sleep stage for each individual. The results were then subjected to group analysis independently for each sleep stage.

For comparisons of the relative band power of 0.2–0.3 Hz for AF between different sleep stages, the statistical analysis was similar to that for BF (see Figures 8C and F for the results). For the significance test of the correlation coefficients, they were first subjected to Fisher z-transformation using the “atanh” function in MATLAB. Then, a one-sample t-test was conducted for the group coefficients that met the normality, whereas the Wilcoxon signed-rank test was conducted for those that did not meet the normality. The tests were performed independently for each sleep stage (see Figure S2 for the results).
